# The repertoire of serous ovarian cancer non-genetic heterogeneity revealed by single-cell sequencing of normal fallopian tube epithelial cells

**DOI:** 10.1101/672626

**Authors:** Zhiyuan Hu, Mara Artibani, Abdulkhaliq Alsaadi, Nina Wietek, Matteo Morotti, Laura Santana Gonzalez, Salma El-Sahhar, Mohammad KaramiNejadRanjbar, Garry Mallett, Tingyan Shi, Kenta Masuda, Yiyan Zheng, Kay Chong, Stephen Damato, Sunanda Dhar, Riccardo Garruto Campanile, Hooman Soleymani majd, Vincenzo Cerundolo, Tatjana Sauka-Spengler, Christopher Yau, Ahmed A. Ahmed

## Abstract

The inter-differentiation between cell states promotes cancer cell survival under stress and fosters non-genetic heterogeneity (NGH). NGH is, therefore, a surrogate of tumor resilience but its quantification is confounded by genetic heterogeneity. Here we show that NGH can be accurately measured when informed by the molecular signatures of the normal cells of origin. We surveyed the transcriptomes of ∼ 4000 normal fallopian tube epithelial (FTE) cells, the cells of origin of serous ovarian cancer (SOC), and identified six FTE subtypes. We used subtype signatures to deconvolute SOC expression data and found substantial intra-tumor NGH that was previously unrecognized. Importantly, NGH-based stratification of ∼1700 tumors robustly predicted survival. Our findings lay the foundation for accurate prognostic and therapeutic stratification of SOC.

**Highlights:**

1. The projection of FTE subtypes refines the molecular classification of serous OC
2. Comprehensive single-cell profiling of FTE cells identifies 6 molecular subtypes
3. Substantial non-genetic heterogeneity of HGSOC identified in 1700 tumors
4. A mesenchymal-high HGSOC subtype is robustly correlated with poor prognosis

## Introduction

Intratumor heterogeneity is a key mechanism for survival and evolution of tumors (Brock et al., 2009; Marusyk et al., 2012). Genetic heterogeneity has been established as a mechanism of survival for most cancer types (Greaves and Maley, 2012; McGranahan and Swanton, 2017; Nowell, 1976). Additionally, cancer cells of the same genetic background may have different phenotypic cell states that enable essential tumor characteristics such as invasion, metastasis and resistance to chemotherapy (Brock et al., 2009; Pisco et al., 2013). The ability of cancer cells to change from one cell state to another (plasticity) is a key feature for cancer survival (Meacham and Morrison, 2013). While genetic heterogeneity is acquired, phenotypic heterogeneity is often inherited from the parent cell-of-origin of a tumor (Gupta et al., 2011; Visvader, 2011). Understanding cancer cell plasticity is dependent on the accurate identification and characterization of individual cell states. Recognizing that cancer cells have plasticity should be reflected on novel tumor stratification strategies that take into account the co-existence of multiple cancer cell states within any one tumor. However, the direct molecular characterization and identification of such states in a tumor is significantly confounded by genetic heterogeneity. An alternative approach for elucidating the phenotypic repertoire of a tumor type is to study the cell states of its cell-of-origin and use those characteristics to decompose an individual tumor into its constituents (Baron et al., 2016; Gupta et al., 2011; Newman et al., 2015).

High-grade serous ovarian carcinoma (HGSOC) is the most aggressive gynecological malignancy (Koshiyama et al., 2017), which is characterized by ubiquitous *TP53* mutations and frequent chromosomal alterations (Ahmed et al., 2010; Bell et al., 2011; Etemadmoghadam et al., 2009; Macintyre et al., 2018). One of the major challenges of HGSOC is the late presentation with almost 80% of patients diagnosed at Stage III or IV disease and a five-year survival rate of about 30% (Torre et al., 2018). The absence of robust methods to stratify HGSOC has been a challenge against therapeutic innovation. Previous studies have demonstrated the transcriptomic heterogeneity of HGSOC (Bell et al., 2011; Tothill et al., 2008). However, analyses using bulk transcriptomes are confounded by a variety of factors, such as copy number variation (Macintyre et al., 2018) and infiltration by non-cancer cells. Importantly, such analyses do not take into consideration the possibility of multiple cell states co-existing in one tumor. Such confounding factors lead to unstable classifications with variable prognostic power across independent datasets (Chen et al., 2018). It is plausible to hypothesize that phenotypic states of the cell-of-origin of ovarian cancer may be echoed in daughter cancer cells. Linking basal and luminal cells from the mammary epithelium to corresponding breast cancer subtypes is a clear example of achieving a stable molecular classification that is based on understanding of phenotypic diversity of the cell-of-origin. This approach has also been successful in other cancers (Fessler and Medema, 2016; Gilbertson, 2011; Ince et al., 2007; Wang et al., 2013).

Recent studies strongly support that serous ovarian cancer can originate from the fallopian tube epithelium (FTE) with evidence from mouse models and genetic evolutionary studies (Ducie et al., 2017; Kim et al., 2012; Labidi-Galy et al., 2017; Perets et al., 2013). Previous work reported that the FTE is composed of PAX8 positive secretory, TUBB4 positive ciliated and CD44 positive basal cells (Clyman, 1966). However, whether there are additional cellular subtypes and whether these subtypes are connected to subtypes of HGSOC have remained elusive.

In this study, for the first time, we analyzed the transcriptomes of around four thousand single cells from normal fallopian tubes from eleven donors and identified six subtypes of FTE cells. By decomposing the expression data from bulk tumors, we matched tumor subtypes with FTE cellular subtypes and revealed a robust prognostic classifier for serous ovarian cancer.

## Results

### Culturing substantially alters single-cell transcriptomes

We analyzed 3,968 single cells from the fallopian tubes of six ovarian cancer patients and five endometrial cancer patients using the Smart-Seq2 technique (Picelli et al., 2014) (Figure 1A and Table S1). Flow Cytometry was used to identify and sort single epithelial cells (EPCAM+, CD45-), leukocytes (EPCAM-, CD45+) and stromal cells (EPCAM-, CD45-) prior to sequencing. To overcome the confounding batch effects and patient-specific variability in clinical samples, we used differential-expression-based clustering (Figure 1B and Methods). Using this approach, we successfully differentiated between epithelial and non-epithelial cells (Figure 1C).

**Figure 1.**
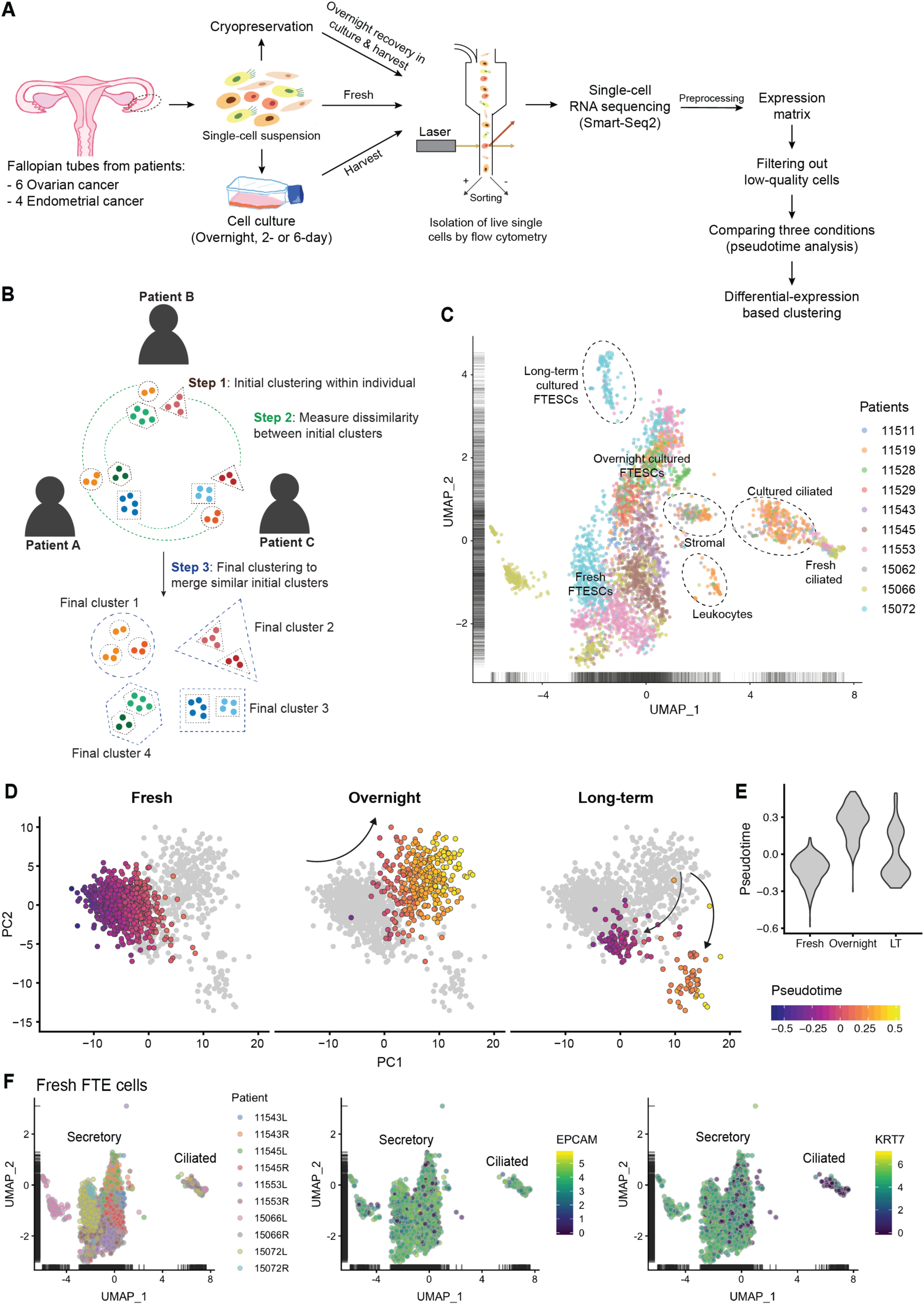
A cell census of human fallopian tubes. (A) A diagram showing the single-cell RNA sequencing and analysis workflow. (B) A diagram showing the clustering approach. Three steps are used to overcome the confounding batch effects and inter-patient variability (see Methods). (C) Uniform manifold approximation and projection (UMAP) plot showing the dimensionality reduction of ∼3,800 single-cell transcriptomes from fallopian tube cells. The cells are colored by their patient sources and annotated with cluster names. (D) Principal component (PC) analysis plot showing the cells under different condition (fresh, overnight-cultured or long-term cultured) colored by the imputed pseudotime values. Pseudotime analysis of secretory cells suggested a more deviated transcriptome after overnight culture compared to the long-term cultured cells. (E) Violin plot showing the distribution of pseudotime under different conditions. The y axis corresponds to the predicted pseudotime and the x axis corresponds to the conditions. Pseudotime analysis here is defined as the quantification of temporal ordering by projecting high-dimensional data to one dimension. LT, long-term cultured. (F) UMAP plots of fresh epithelial cells showing the difference between secretory cells and ciliated cells. Each dot denotes one cell. In the left panel, the cells are colored by their donors and the side of tubes (L, left; R, right); while they are colored by the expression levels of an epithelial marker, *EPCAM*, in the middle panel or the expression levels of a secretory marker, *KRT7*, in the right panel as shown in the scale bar.

However, we observed striking effects induced by culture conditions on the transcriptomes of single cells by comparing freshly dissociated cells with cultured ones (Figures S1A-D and Tables S2, S3). For example, overnight culture induced the expression of genes that are rarely expressed in FTE cells such as *CD44* (log_2_ fold-change [log-FC] = 3.8) (Paik et al., 2012) and reduced the expression of key markers of secretory cells, such as Estrogen Receptor alpha (*ESR1*) and Oviductal Glycoprotein 1 (*OVGP1*) (Figure S1E and Table S3) (Cerny et al., 2016; Wu et al., 2016). Cilium organization was also downregulated in the overnight-cultured ciliated cells. Moreover, the Pseudotime analysis (Campbell and Yau, 2018) across three conditions revealed that the transcriptomes of long-term (LT) cultured cells were more similar to the fresh cells compared to the overnight-cultured group (Figures 1D and 1E). For instance, the fatty acid metabolic process (e.g. *BDH2*, *ALKBH7* and *PTGR1*) was transiently downregulated after overnight culture and then upregulated in the LT group, while the RNA processing pathway was upregulated temporarily (Figures S1A-B and S1F). This suggests that including the overnight-cultured cells in subsequent analysis may introduce significant biases that would preclude meaningful conclusions. Similarly, although the LT group resembled the fresh group of cells, they showed a unique split into two sub-groups and perturbed expression of Stathmin (*STMN1*) as well as cell cycle genes that probably represent an artefact of long-term culture (Figures S1A and S1G). Therefore, overnight and long-term culture likely introduced nontrivial alteration in gene expression. To avoid these substantial effects from preservation methods, we focused our downstream analysis on fresh cells only. Although fresh cells were affected by early response genes (e.g. *FOS* and *JUN*), our analysis indicates that such affected cells can be detected by unsupervised clustering.

### A cell census of human fallopian tubes in cancer patients

Within epithelial cells, we identified two previously established subtypes, secretory and ciliated cells (Figure 1F). Secretory cells were characterized by the expression of *PAX8* and *KRT7* (Figure S1H) as well as a large number of newly identified markers of secretory cells (Table S4). The ciliated population was represented by the strong expression of *FOXJ1* and members of the coiled-coil domain containing protein family, such as *CCDC17* and *CCDC78* (Figures S1I and J). This protein family is essential for cilia functioning (Klos Dehring et al., 2013). We also identified a list of previously unrecognized markers of fallopian tube ciliated cells (Table S4), such as the calcium binding protein Calcyphosin (*CAPS*), that were enriched in the cilium-related pathways (Figures S1I and S1K) (Wang et al., 2002).

We next identified secretory cell subtypes based on their transcriptomes. Only fresh secretory cells with strong expression of *KRT7* and *EPCAM* and no expression of *CCDC17* or *PTPRC* (also known as CD45) were included in the analysis. In addition, to avoid including contaminating cancer cells, we excluded cells that had detectable copy number variants or apparent loss-of-heterozgyosity (Figure S2A) (Fan et al., 2018). We applied the aforementioned differential-expression-based clustering method and identified nine clusters with distinct transcriptional profiles within the secretory cell population (Figures 2A and 2B). Except for a patient-specific cluster (C8) that was enriched in inflammatory markers, all other clusters contained cells from multiple patients (Figure 2A). Cluster 8 was, therefore, not considered for further analysis. Three out of nine clusters (C1, C2 and C5) had no particular distinguishing features and clusters C1 and C2 had low library complexity (Figure S2B). This result suggests that these three clusters probably represented a quiescent population due to cell senescence or loss of hormonal influence (CROW et al., 1994; van Deursen, 2014). These clusters were, therefore, excluded from further analysis. Cluster C6 had evidence of cell stress as shown by the high expression levels of early response genes, such as *FOS* and *JUN* (Honkaniemi et al., 1992). This cluster was excluded from further analysis because such stress response is probably the result of sample preparation prior to sequencing.

**Figure 2.**
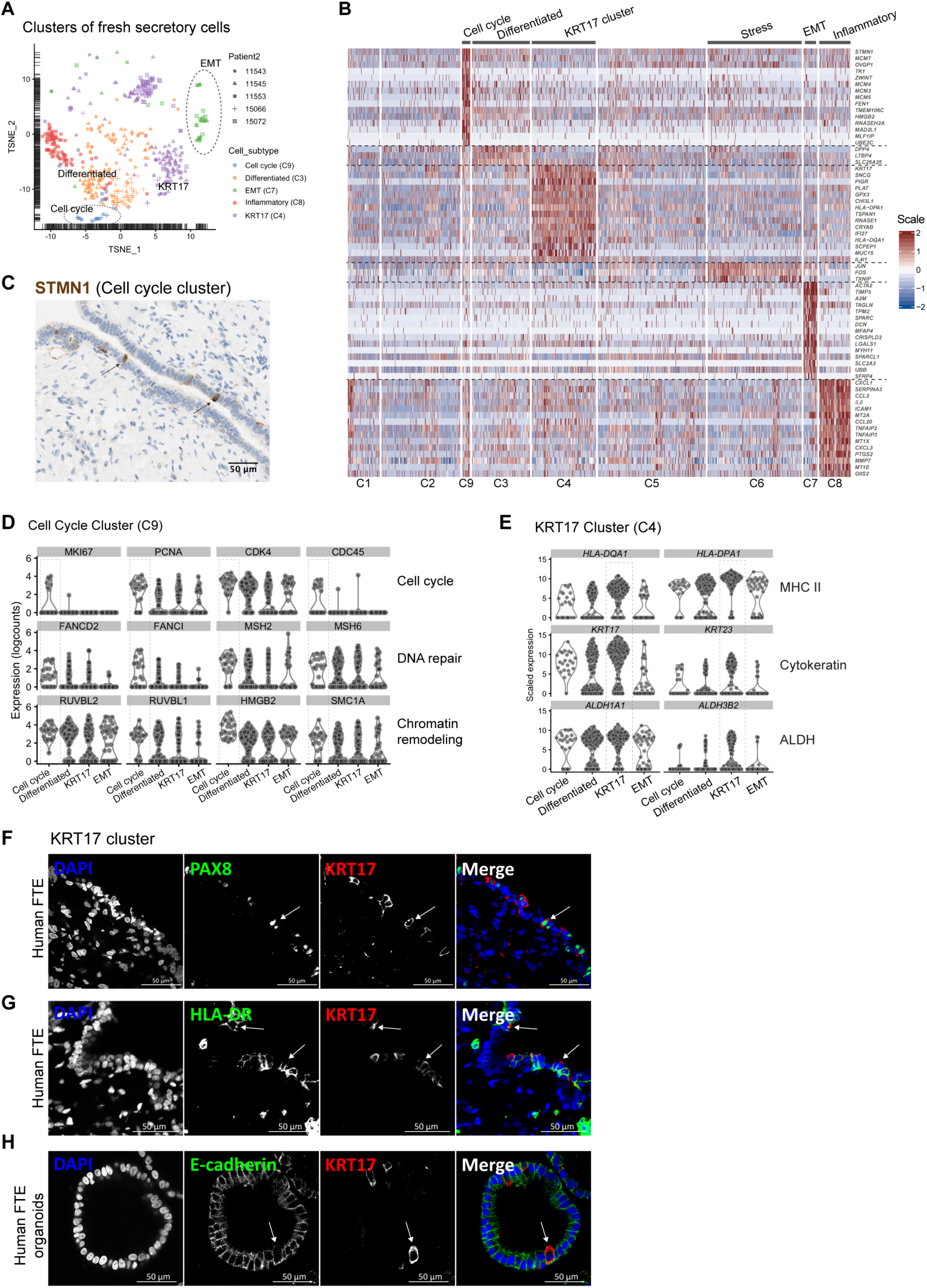
Single-cell RNA sequencing of FTE cells identifies four novel secretory subtypes. (A) t-Distributed Stochastic Neighbor Embedding (t-SNE) plot showing the clusters within fresh secretory cells. Each point represents one cell that is colored by its cluster/subtype and shaped by its donor, as shown in the legend. The quiescent population and the stress population are not shown in this plot. Apart from C8, other clusters are composed of cells from multiple donors. (B) Heatmap showing the clustering result of fresh secretory cells and the expression levels of their top marker genes. Each column represents a single cell and each row represents a top marker gene. The color scale indicates the expression level shown in the scale bar. Based on this clustering result, we identified four novel secretory cell subtypes that are termed cell cycle (C9), differentiated (C3), KRT17 cluster (C4) and EMT (C7). C3 represents cells that are under stress and C8 is a patient-specific inflammatory cell population. (C) IHC staining confirming the existence of the cell cycle cluster (arrow) by its marker *STMN1* (Stathmin) in a human fallopian tube section. (D) Violin plots showing the upregulation of nine representative marker genes of the Cell Cycle Cluster (C9) that are respectively related to the cell cycle, DNA repair or chromatin remodeling pathways. (E) Violin plots showing the upregulation of six representative marker genes of the KRT17 cluster (C4) that respectively belong to MHCII, cytokeratins or aldehyde dehydrogenases (ALDH). (F-G) IF staining validating the KRT17 cluster by using the antibodies against KRT17, a secretory marker PAX8 and HLA-DR in the human fallopian tube sections. (F) demonstrates that KRT17 positive cells are also PAX8 positive, while (G) confirms the co-expression of HLA-DR and KRT17 in this subpopulation. (H) IF double staining of KRT17 and an epithelial marker E-cadherin in the organoid derived from human FTE. It indicates that this KRT17 cluster is a stable population that can be derived *ex-vivo* using organoid cultures.

Cluster C9, that we termed cell cycle cluster, comprised ∼1.6% of fresh FTESCs and was enriched in three pathways, namely cell cycle (e.g. *MCM2-7*, *MKI67*, *TK1* and *STMN1*), DNA repair (e.g. *FANCD2*, *FANCI* and *MSH2*) and chromatin remodeling (e.g. *HMGB2* and *SMC1A*) (Figures 2B-D). *MKI67* (also known as Ki-67) is a well-known marker for proliferation in FTE and other cells (Kuhn et al., 2012). The two Fanconi Anemia proteins, FANCD2 and FANCI, can form a heterodimer that is essential for DNA repair and can interact with MCM2-7 (Nalepa and Clapp, 2018). The relatively low percentage of cycling cells is consistent with the ages and postmenopausal status of patients from whom the cells were obtained. Cluster C3, was termed the differentiated subtype, had significantly increased representation from genes involved in RNA synthesis and transport pathways (e.g. *PTBP1*, *ZNF259* and *PRPF38A*). It also shared several markers with cluster C9. This may represent a transient differentiating cell population following cell division.

Cluster C4, termed the KRT17 subtype, was characterized by the upregulated expression of major histocompatibility complex (MHC) Class II genes (e.g. *HLA-DQA1*, *HLA-DPA1* and *HLA-DPB1*), cytokeratins (*KRT17* and *KRT23*), aldehyde dehydrogenases (e.g. *ALDH1A1* and *ALDH3B2*) and *CDKN1A* (also called p21) (Figures 2B and 2E, Table S5). This subpopulation of secretory FTE cells that are enriched in MHC Class II expression has not been characterized previously (Comer et al., 1998). Importantly, we validated this KRT17 positive cluster in human FTE and organoids derived from human FTE cells, suggesting that it represents a robust group of cells with potentially important biological functions (Figures 2F-H).

C7 showed high expression of a regulator of G protein signaling (*RGS16*) and genes enriched in the extracellular matrix (ECM) pathway (false discovery rate [FDR] = 1.80e-17), such as *TIMP3* and *SPARC* (Figures 2B, 3A, B and Table S5). We assumed that this cell type is generated by partial epithelial-mesenchymal transition (EMT) (Nieto et al., 2016), which can be induced by the chronic exposure to oxidative stress (Mahalingaiah et al., 2015) and contribute to cancer development (Hanahan and Weinberg, 2011). Hence, we termed C7 as the EMT subtype. To verify that this cell type is not a contamination from FT stromal cells, we checked the expression level of epithelial markers, doublet likelihood and the expression of stromal markers. This EMT population strongly expressed *KRT7* and *EPCAM* (Figure 3A) as expected following the aforementioned filtering. It also passed our filter for doublets; 40 (100%) out of 40 cells in the EMT cluster were detected as singlets (McGinnis et al., 2019). Moreover, two stromal markers (*COL1A2* and *COL3A1*) were specifically expressed in stromal cells but not in the EMT population (Figure S3A). These results demonstrate that this population is not contaminated by mesenchymal cells.

**Figure 3.**
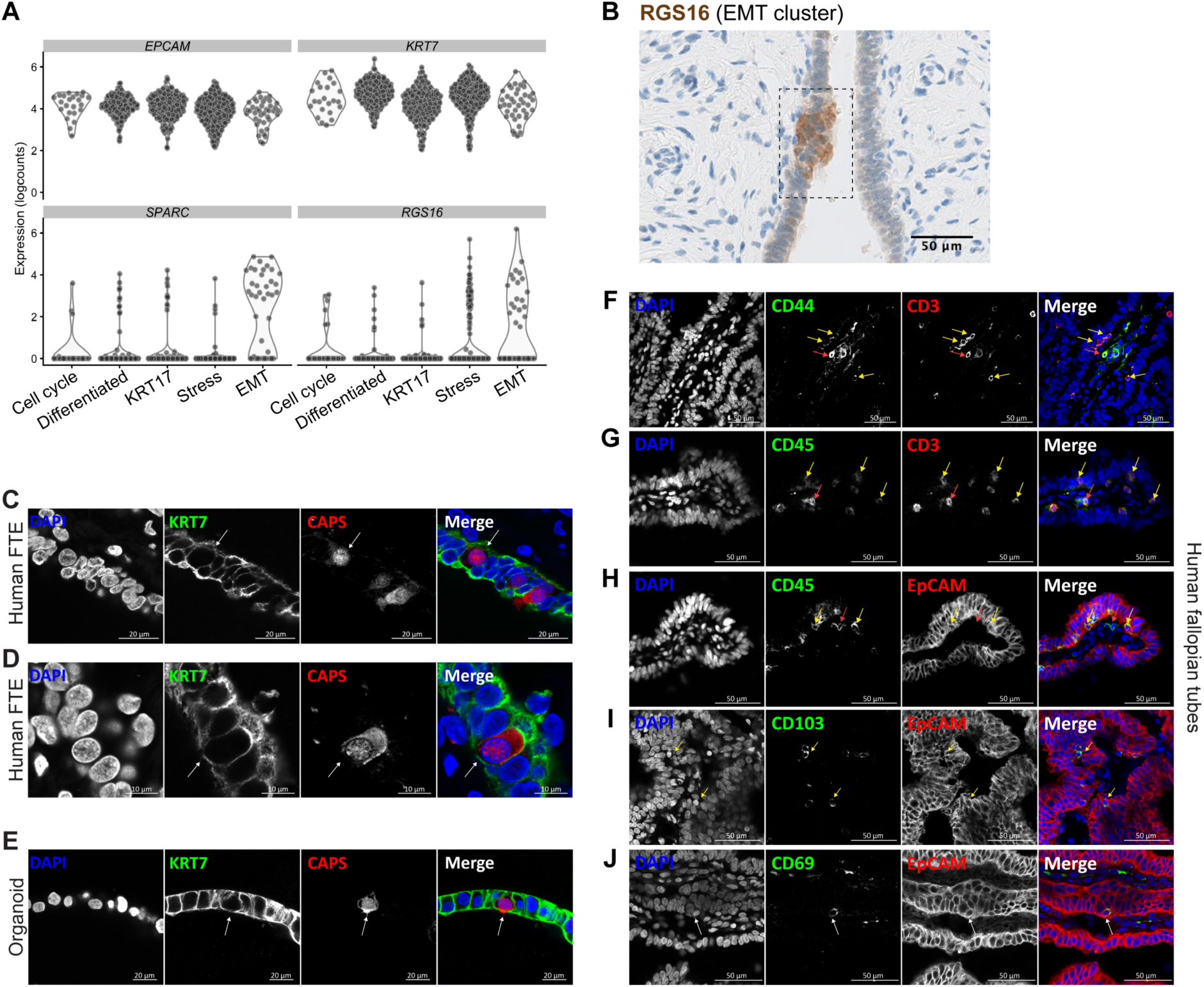
Non-traditional cell subtypes in the fallopian tube epithelial layer. (A) Violin plots showing that the EMT cluster of FTE cells has strong expression of *KRT7* and *EPCAM* as well as the upregulated expression of *SPARC* and *RGS16*. (B) IHC staining validating the EMT Cluster with its marker, RGS16, in human FTE (dashed box). (C-D) IF staining showing the KRT7 and CAPS double positive intermediate cell population (arrows) in the human FTE. (E) IF staining showing an example of the KRT7+CAPS+ intermediate cell population (arrow) in the human fallopian tube organoid cultured. (F-H) IF staining indicating the EPCAM+ CD44+ presumed “peg cells” that are positive for lymphocyte markers CD45 and CD3 as shown by co-expression of CD44 and CD3 (F), CD45 and CD3 (G) as well as CD45 and EpCAM (H) in the human FTE. The intra-epithelial CD44+ CD3+ cells are also CD45+ and EpCAM+ (yellow arrows). This indicates that the basal CD44+ cells are T lymphocytes. We also observe the extra-epithelial CD44+ CD3+ CD45+ EpCAM-cells in the stromal region (red arrows). (I-J) IF staining showing that the basal cells are positive for EPCAM and two markers of tissue-resident memory T lymphocytes, CD103 (I) and CD69 (J), in the human FTE.

In addition to the four secretory cell types, we discovered a rare intermediate cell type that was characterized by the expression of the secretory cell marker KRT7 and high expression of the ciliated cell marker CAPS (Figures 3C-D and S3B-G), whilst other KRT7 positive secretory cells were CAPS negative. *PAX8* was expressed in a subset of this intermediate population, possibly due to its moderate expression level leading to a higher dropout rate. This population was validated in human FTE tissue sections (Figures 3C and 3D). Additionally, this subtype was enriched in overnight cultured cells and recapitulated in the three-dimensional organoid culture derived from human FTE tissues (Figure 3E). This intermediate population most probably represents an intermediate state between secretory and ciliated cells, which accords with the previously assumed transition from secretory to ciliated cells (Ghosh et al., 2017; Hellner et al., 2016).

Because we observed that chemokines and MHC genes were frequently expressed in FTE cells (Figure 2B), we examined fallopian tube sections for the expression of markers of lymphocytes and identified a basal CD45+EPCAM+ cell population. This population was also positive for CD3, CD44, CD69 and CD103 (Figures 3F-J), suggesting that these basal cells are tissue-resident memory T lymphocytes (TRMs) (Topham and Reilly, 2018) and that the FTE is not immunologically inert. This is in line with previous studies (Ardighieri et al., 2014; Peters, 1986). Nevertheless, we demonstrate that this basal cell population is also EPCAM positive, which is surprising because TRM lymphocytes have not been previously reported to be positive for epithelial markers in human.

### Revealing the cell state composition of HGSOCs using deconvolution

Given the link between types of cell of origin and tumor types that has been reported in other cancers such as breast cancer (Bertucci et al., 2012; Dai et al., 2015) and the known shared key markers between HGSOC and FTE cells (such as *PAX8*, *WT1* and *ESR1*) (Ince et al., 2015; Perets et al., 2013), we hypothesized that HGSOC cell states are linked to FTE cell subtypes. Based on our profiling of single-cell transcriptomes, we computed a reference matrix with cell-type derived transcriptomic signatures from the five major FTE cellular subtypes (cell cycle, EMT, differentiated, KRT17 cluster and ciliated) as previously described (Baron et al., 2016) (Figure 4A and Table S6). The resulting reference matrix was then used in the deconvolution analysis (Newman et al., 2015) of the bulk HGSOC RNA-seq data from The Cancer Genome Atlas (TCGA) (Bell et al., 2011) and the microarray data from the Australian Ovarian Cancer Study (AOCS) (Tothill et al., 2008) to compute the factions of five cell states within each tumor. The decomposition of bulk tumor samples revealed intra-tumor transcriptomic heterogeneity according to the proportions of cell subtypes in both datasets (Figures 4A-E). Remarkably, the ciliated tumor subtype was highly enriched in the low-grade tumors (Grade 1) compared to the high-grade ones (Grades 2 and 3) in the AOCS dataset and an additional dataset that also comprised both high-grade and low-grade tumors (Yoshihara et al., 2010) (p = 1.3e-10, one-sided Wilcox test, Figures 4E, F and S4A). This strongly suggests that grades of serous ovarian carcinoma are associated with their ability to differentiate into cells that molecularly resemble FTE ciliated cells. In contrast, the TCGA dataset, which only included HGSOCs, had no tumors that were enriched in the ciliated subtype (Figure 4B). Therefore, the enrichment in these ciliated markers is most probably a distinguishing molecular feature of low-grade tumors.

**Figure 4.**
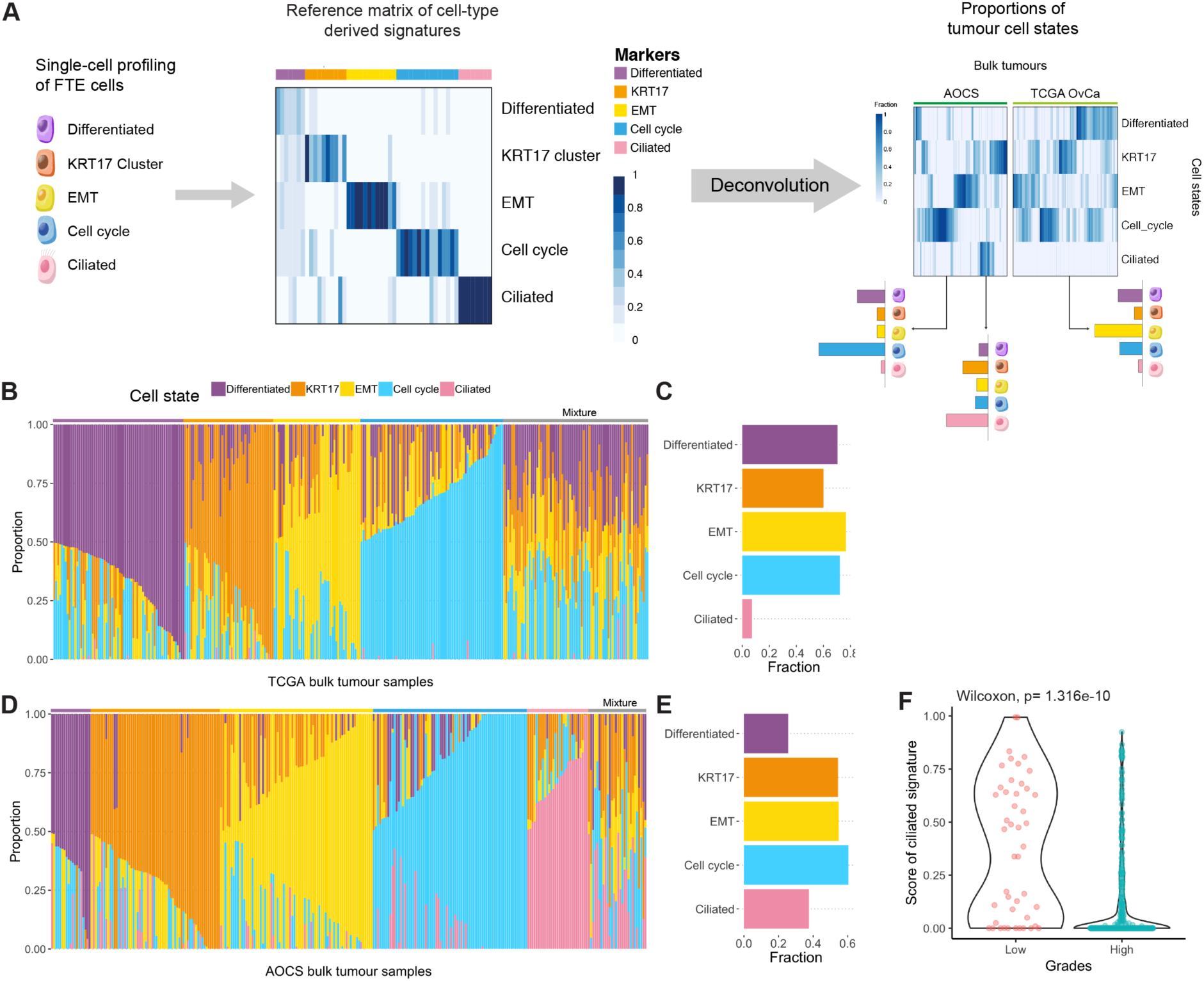
The repertoire of phenotypic heterogeneity of serous ovarian cancer revealed by deconvolution analysis. (A) A reference was calculated from our scRNA-seq data of FTE cells, where columns correspond to five cell subtypes in FTE as indicated. The heatmap titled the reference matrix depicts the magnitude of the expression levels of the 52 marker genes (rows) across five cell subtypes (see also Table S6). The heatmaps in the right panel depicts the proportions of cell states (rows) across bulk tumor samples (columns) in AOCS and TCGA datasets. Beneath these heatmaps, three cartoon barplots show the schematic composition of three tumor samples. (B) Stacked barplot showing the deconvolution result of 308 tumors from the TCGA ovarian carcinoma study. Colors of the bar denote five cell states as shown in the legend. The y axis represents the proportion of each state across bulk tumor samples (x axis). The annotation bar above the barplot denotes the subtypes of bulk tumors that are defined by the dominant cell state within each tumor, where grey represents the mixture of multiple cell states. (C) Barplot showing the prevalence of five cell states in the TCGA dataset. The x axis represents the proportion of tumor samples in TCGA that harbor the given cell state (y axis). (D) Stacked barplot showing the deconvolution result of 285 tumors from the AOCS dataset as in B. (E) Barplot showing the prevalence of five cell states in the AOCS dataset. Note that the ciliated tumor subtype is absent from the TCGA cohort, which exclusively comprises high-grade tumors. In contrast, the ciliated cell state is more prevalent in the AOCS dataset that contains both low-grade and high-grade tumors. (F) Violin plot showing the significant upregulation of ciliated scores in low-grade serous ovarian tumors compared to high-grade ones. We combined samples from both AOCS (low-grade, n = 20; high grade, n = 260) (Tothill et al., 2008) and GSE17260 (low-grade, 26; high grade, 84) (Yoshihara et al., 2010) that have more than 20 low-grade serous tumors per dataset from CuratedOvarianData. Each dot represents one sample, and it is colored by the grade of serous ovarian cancer. The overall p-value is 1.316e-10, by one-sided Wilcoxon rank sum test.

Most notably, we identified a class of EMT-enriched tumors in multiple datasets. These tumors were enriched in the genes previously linked to the “mesenchymal” HGSOC subtype. We found that the marker genes of these tumors were enriched in the extracellular matrix, focal adhesion and PI3K-Akt signaling pathways (FDR < 0.0002, by DAVID, Table S7), that are critical for tumor cell survival (Fresno Vara et al., 2004; McLean et al., 2005). Furthermore, three key EMT transcription factors, *TWIST1*, *TWIST2* and *SNAI2* (Ansieau et al., 2008; Kang and Massagué, 2004; Yang et al., 2004), were upregulated in the EMT-high tumors (Figure 5A), while the miRNA-200 family (miR-200a, miR-200b, miR-200c, miR-141 and miR-429) that represses EMT (Gregory et al., 2008; Nieto et al., 2016) was downregulated in EMT-high tumors (FDR < 0.01, log-FC < −0.5, Figure 5B), suggesting that EMT may be the underlying mechanism of enrichment of mesenchymal cancer cells in this tumor subtype. We also found that miRNA-483 and miRNA-214 were significantly upregulated in EMT-high tumors, while miRNA-513c, miRNA-509 and miRNA-514 were downregulated (Figure 5B). Although previous studies suggested that miRNA-483 and miRNA-214 play an important role in cancer progression (Chandrasekaran et al., 2016; Liu et al., 2013), their regulatory role in the EMT process has not been evaluated.

**Figure 5.**
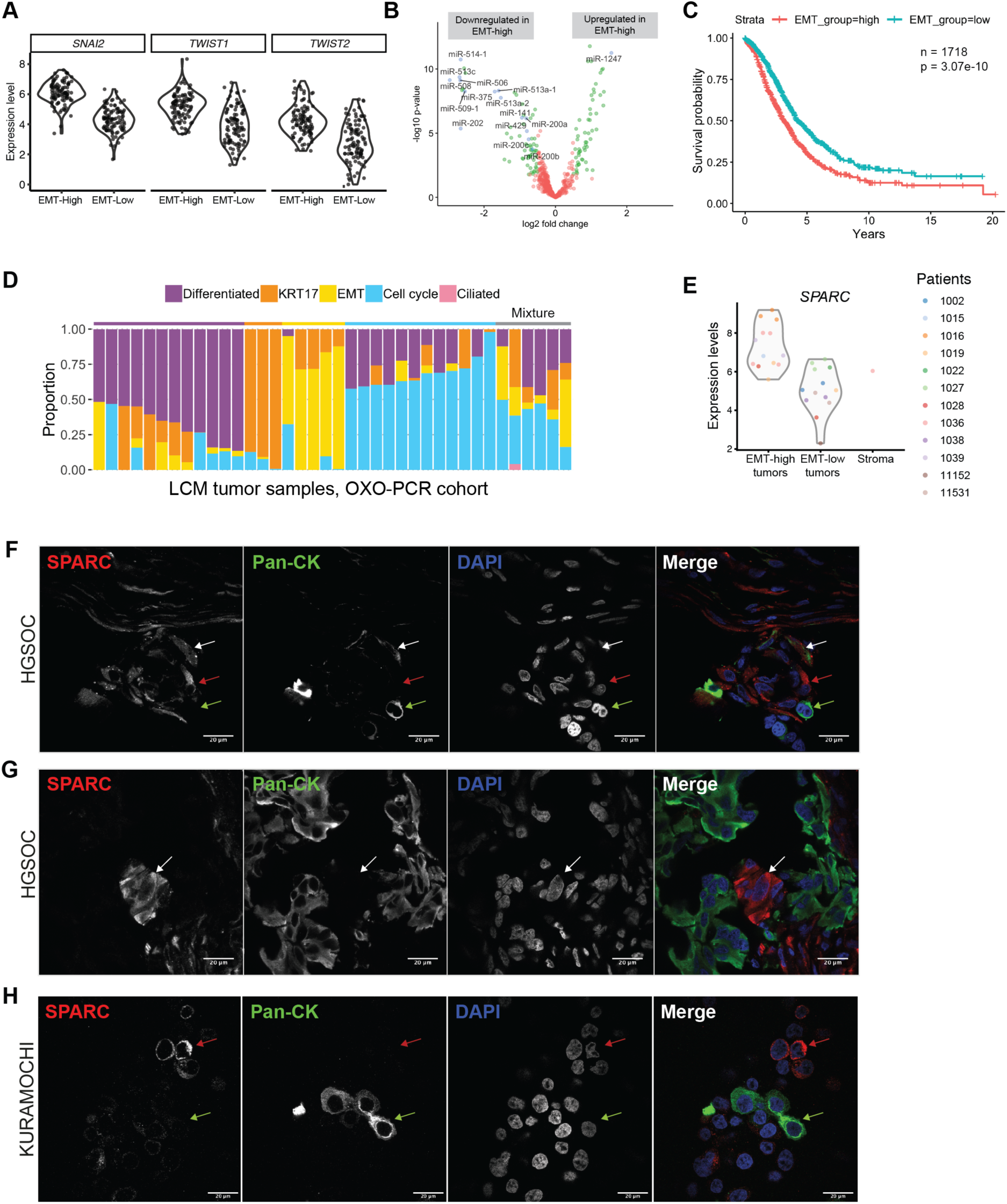
EMT-high tumors comprise mesenchymal-like cancer cells. (A) Violin plots showing the key EMT drivers, Twist (*TWIST1* and *TWIST2*) and Snail (*SNAI2*), that are significantly upregulated in in the EMT-high tumors compared to EMT-low ones in the TCGA dataset (log-FC > 1.8, FDR < 2e-14, by limma voom). (B) Volcano plot showing the miRNAs that are differentially expressed between EMT-high and EMT-low tumors in the TCGA dataset, including the miRNA-200 family (miR-200a, miR-200b, miR-200c, miR-141 and miR-429), which are the suppressors of EMT. The green and blue dots correspond to the miRNAs that are significantly differentially expressed (log-FC > 0.5, FDR < 0.05). (C) Kaplan-Meier plot showing the effect of high EMT scores on survival. The cases (n = 1,731) are combined from nine datasets (see Table 1). Patients are dichotomized into EMT-high (red) and EMT-low (turquoise) by the median of EMT scores in each dataset. Each short vertical line indicates a censoring event. P value = 2.81e-10, by log-rank test, HR = 1.5, CI = 1.3 – 1.7. (D) Stacked barplot visualizing the deconvolution result of 36 LCM tumor samples collected from 15 patients with HGSOC in the OXO-PCR cohort. The y axis denotes the proportion (0-1) of the five cell states (colors) across LCM tumor samples (rows). The annotation bar above the barplot corresponds to the tumor subtypes that are defined by the dominant cell state within each tumor, where grey represents the mixture of multiple cell states (see also Figure 4B). LCM, laser capture microdissection. (E) Violin plot showing the expression levels of the EMT marker that is upregulated in the EMT-high LCM tumor samples compared to the EMT-low ones with one LCM stromal sample as control (p < 0.001, one-sided Wilcox test). Each dot represents a sample colored by the patient ID. (F) IF staining showing the mesenchymal-like tumor cells in the pre-chemotherapy tumor frozen section, by using an EMT marker, SPARC, and an epithelial marker, pan-cytokeratin. The mesenchymal-like tumor cells are double positive for SPARC and pan-cytokeratin (white arrow), while other cells are SPARC positive only (red arrow) or pan-cytokeratin positive only (yellow arrow). Pan-CK, pan-cytokeratin. (G) IF staining showing a mesenchymal-like tumor cell that is SPARC positive and pan-cytokeratin negative (arrow) and has the similar nuclear morphology as the surrounding tumor cells in the human tumor frozen section. (H) IF staining showing mesenchymal-like tumor cells in the KURAMOCHI cell line. There are at least two phenotypes. The mesenchymal-like phenotype is SPARC positive and pan-cytokeratin negative (red arrow), while the epithelial phenotype is SPARC negative and pan-cytokeratin positive (green arrow).

**Table 1.** EMT-high tumors are robustly correlated with poor prognosis.

| GEO dataset | Samples | Hazard ratio | P-value | 95% confidence interval | Citation |
| --- | --- | --- | --- | --- | --- |
| **E.MTAB.386** | 129 | 3.31 | 0.0120 | 1.30 - 8.41 | (Bentink et al., 2012) |
| **GSE49997** | 194 | 3.08 | 0.0038 | 1.44 - 6.60 | (Pils et al., 2012) |
| **GSE13876** | 144 | 2.58 | 0.0272 | 1.06 – 5.11 | (Crijns et al., 2009) |
| **GSE26712** | 185 | 2.03 | 0.0315 | 1.07 – 3.89 | (Bonome et al., 2008) |
| **GSE26193** | 107 | 2.33 | 0.0348 | 1.06 - 5.11 | (Mateescu et al., 2011) |
| **GSE51088** | 139 | 1.89 | 0.0285 | 1.07 - 3.33 | (Karlan et al., 2014) |
| **GSE32062.GPL6480** | 260 | 1.88 | 0.0556 | 0.98 - 3.58 | (Yoshihara et al., 2012) |
| **AOCS (Grade≥2)** | 253 | 2.69 | 0.0004 | 1.56 - 4.66 | (Tothill et al., 2008) |
| **TCGA RNA-seq** | 307 | 2.30 | 0.0047 | 1.29 - 4.09 | (Bell et al., 2011) |

Note: related to Table S8

### EMT-high subtype is robustly correlated with poor prognosis

We next tested whether any of the five tumor subtype scores from the deconvolution analysis was associated with survival. We found that the EMT score was significantly associated with poor overall survival and that this association was independent from the effect of age, stage or residual disease (p < 0.05, by Cox proportional hazard model, Table 1). The robustness of this association was confirmed by permutation testing (n=500) leaving out 10% of the samples each time (empirical p-values = 0.012 [TCGA] and 0 [AOCS], by permutation test). The mesenchymal subtype was previously reported to be associated with poor prognosis (Konecny et al., 2014; Tan et al., 2013), but the reproducibility of this observation was inconsistent probably because of the difficulty in defining this group of tumors. Using EMT scores from deconvolution, we reached a robust classification with consistently significant correlation with poor survival (p < 0.05) in eight independent datasets, including TCGA dataset, AOCS dataset and six additional microarray datasets (n > 100 patients in each set) from the CuratedOvarianData database (Ganzfried et al., 2013) (Table 1). When we combined and dichotomized all the samples (n = 1,718), the EMT-high tumors had significantly worse prognosis (p = 3.07e-10, Figure 5C). This demonstrates that deconvolution analysis using the identified panel of genes is strongly predictive of prognosis in serous ovarian cancer.

To test whether the EMT-high tumor subtype was merely a reflection of stromal cell impurities in tumor samples, we performed RNA sequencing on 36 laser capture microdissected (LCM) tumor samples collected from 15 patients with HGSOC and classified tumors based on the deconvolution analysis (Figure 5D). We next compared the expression of genes in the EMT signature between laser capture microdissected tumor and stroma samples. As expected, the expression levels of *PAX8* and *EPCAM* were significantly higher in both EMT-high and EMT-low tumor samples compared to stroma in which these markers showed almost no expression (Figure S5A). In contrast, the EMT marker was highly expressed in EMT-high tumor confirming that EMT-high tumors truly express these genes (Figure 5E), further confirming EMT-high as a genuine tumor subtype. *SPARC*, one of the EMT markers, was previously described in the mesenchymal subtype of HGSOC (Tothill et al., 2008), while certain other markers were reported to be related to EMT in ovarian cancer or other cancers (Anastassiou et al., 2011; Ford et al., 2013; Li and Yang, 2016). Nevertheless, the link between this tumor subtype and a particular FTE cellular subtype was previously unrevealed. To further confirm that the EMT-high marker SPARC is strongly expressed in cancer cells, we performed immunofluorescent staining on pre-chemotherapy tumor frozen sections, where we indicated cancer cells by co-staining with an antibody that recognized pan-cytokeratin. Our results revealed the co-existence of at least two populations of cancer cells, a pan-cytokeratin high, SPARC negative population and a pan-cytokeratin low and SPARC positive population within the same tumor (Figures 5F, S5B and Supplementary Video). In addition, pan-cytokeratin negative and SPARC high cancer cells could also be identified by their abnormal nuclei (Figures 5G and S5C), which were distinct from the SPARC high fibroblasts cells (Tan et al., 1999). Similarly, this SPARC positive, pan-cytokeratin negative and mesenchymal-like subpopulation was observed in the KURAMOCHI cell line, which is known to faithfully represent HGSOC (Domcke et al., 2013). KURAMOCHI cells exhibited two phenotypes, a mesenchymal-like phenotype that was SPARC positive and pan-cytokeratin negative and an epithelial phenotype that was SPARC negative and pan-cytokeratin positive (Figures 5H and S5D-E), suggesting that this cell line could be a potential *in vitro* model to study the mesenchymal subtype of HGSOC. Importantly, TP53 sequencing of this cell line showed a single peak for the previously described mutation (c.841G>T) in this cell line (Anglesio et al., 2013), validating that the phenotypic heterogeneity was not caused by contamination with another cell line.

## Discussion

In this study, we performed deep single-cell RNA-seq on thousands of nonmalignant epithelial cells from the FTE and uncovered a strong link between diverse secretory cell populations and HGSOC subtypes. This elucidates the repertoire of phenotypic heterogeneity in cancer cells that are inherited from the cell-of-origin. We propose a model whereby individual HGSOCs are composed of a mixture of cancer cell states that we revealed by single-cell sequencing of the putative cell-of-origin (Figure 6). Unraveling the trajectory of differentiation from one cancer cell state to another will be needed in future studies to understand the mechanistic basis of HGSOC cell plasticity.

**Figure 6.**
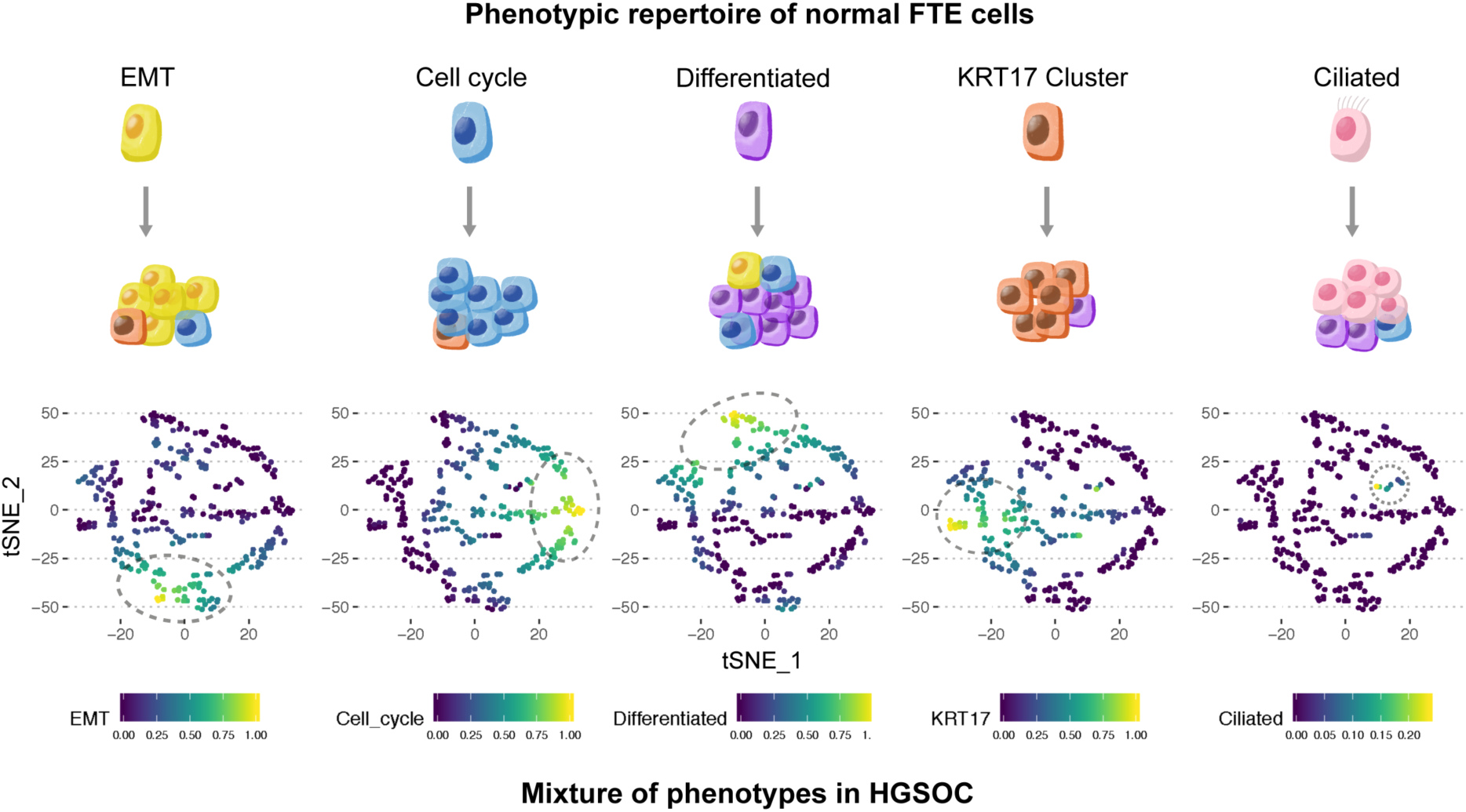
HGSOCs are composed of a mixture of cancer cell states inherited from its cell-of-origin in the FTE. This diagram proposes a model that HGSOCs are assembled from multiple cancer cell states that echo the cellular phenotypes of its tissue-of-origin. The upper panel exhibits five cellular subtypes of FTE. In the lower panel, five t-SNE plots demonstrate the cell state hubs within the deconvolution result of TCGA data (see also Figure 4B). Each dot is a HGSOC bulk sample. The x and y axes represent the first two components of tSNE. The color scale represents the proportion of cancer cell states. Grey dashed circles highlight the hubs representing the tumor subtypes of serous ovarian cancer.

Within the FTE, our analyses recognized a KRT17 positive population with high expression of aldehyde dehydrogenases (ALDHs) and MHCII. ALDH was reported as the marker of mammary stem cells (Biton et al., 2018; Ginestier et al., 2007). Moreover, a subpopulation of intestinal stem cells was recently reported to express MHCII. The latter was found to regulate the stem cell fate through the interaction between T helper cells and stem cells (Biton et al., 2018; Ginestier et al., 2007). Such evidence implies that the KRT17 cluster that we identified could include a progenitor population. This opens a door for studying self-maintenance and differentiation of epithelial cells in fallopian tubes.

Phenotypic diversity exists in HGSOC at the transcriptomic level (Bell et al., 2011; Tothill et al., 2008). Based on the single-cell profiling of FTE cells, our deconvolution analysis matched the FTE cellular subtypes with ovarian cancer cell states within individual tumors. The correspondence between tumors and normal tissues can be explained by at least two hypotheses that are not necessarily mutually exclusive. The first one is that each tumor subtype originates from an individual cell type of origin (Ince et al., 2007; Visvader, 2011). The second hypothesis is that multiple tumor subtypes come from one single cellular subtype, sometimes referred to as the cancer stem cell, while the plasticity of cancer cells mimics that of the normal cell-of-origin (Gupta et al., 2011; Visvader, 2011). Both concepts are compatible with our findings. Lineage tracing will be needed to investigate the underlying mechanism of the correspondence between tumors and the tissue-of-origin.

Despite the fact that the mesenchymal-like tumor subtype was previously reported as a distinct subtype in HGSOC (Tothill et al., 2008), it has been controversial whether the expression of EMT markers originated from tumor cells, infiltrating stromal cells or both (Schwede et al., 2018; Zhang et al., 2019). Our results demonstrate that the expression of mesenchymal-related genes, such as *SPARC*, occurs in tumor cells. This further supports the existence of the EMT process in human tumors, which may enhance their invasion ability and drug resistance (Schmidt et al., 2015; Singh and Settleman, 2010). This notion is consistent with our finding that the EMT-high HGSOC subtype is related to poor survival. However, we cannot exclude the possibility that the expression of EMT genes in bulk tumor analyses may arise from stromal infiltration or cancer-associated fibroblasts (CAFs). Since CAFs can potentially promote cancer progression (G. M. Chen et al., 2018; Kalluri, 2016; Schwede et al., 2018; (Eckert et al., 2019), it will also be important to scrutinize the role of CAFs in HGSOC and the interaction between CAFs and ovarian cancer cells.

Compared to the previously developed prognostic signatures of HGSOC that were difficult to be reproduced in multiple datasets (Bell et al., 2011; Bonome et al., 2008; Chen et al., 2018; Jiang et al., 2006; Karlan et al., 2014; Konecny et al., 2014), our classifier is strongly reproducible across eight independent datasets and is based on the expression of only fifty-two marker genes. Robust identification of tumor subtypes is a prerequisite for identifying treatment strategies that limit the ability of cancer cell to acquire a mesenchymal state (Davis et al., 2014; Ishay-Ronen et al., 2019). Given these factors, our approach possesses potential translational significance to improve the survival of HGSOC patients.

In conclusion, our single-cell profiling illuminates the cellular landscape of human fallopian tube epithelium, the presumed tissue-of-origin of HGSOC, for the first time. Our work not only expands the understanding of FTE, but also provides the first benchmark dataset on FTE to study HGSOC. The combined analysis of single-cell data and tumor expression data revealed the connection between FTE cellular subtypes and HGSOC tumor subtypes and identified the EMT-high subtype with important clinical implications.

## Methods

### EXPERIMENTAL SUBJECT DETAILS

#### Cell line and cell culture

KUARMOCHI cell line (cell number ID: JCRB0098) was obtained from Japanese Collection of Research Bioresources (JCRB) Cell Bank. KURAMOCHI cells were cultured in RPMI 1640 media (Gibco) with Fetal Bovine Serum (FBS, 10%; Gibco) and Penicillin-Streptomycin (100 U; Gibco) at 37°C, 5% CO_2_ and 95% humidity.

#### Human fallopian tube and tumor samples

The cases in this study were recruited under the Gynecological Oncology Targeted Therapy Study 01 (GO-Target-01, research ethics approval #11-SC-0014) and the Oxford Ovarian Cancer Predict Chemotherapy Response Trial (OXO-PCR-01, research ethics approval #12-SC-0404). All participants involved in this study were appropriately informed and consented. Fallopian tube (FT) biopsies for single-cell RNA-seq were collected from the distal end of fallopian tubes. In total, we collected FT samples from six patients with HGSOC, five with endometrial cancer (Table S1). Tumor samples were biopsied during diagnostic laparoscopy, immediately frozen on dry ice and stored in clearly labelled cryovials in −80 °C freezers.

### METHOD DETAILS

#### Sample dissociation, culture and cryopreservation

The fallopian tube samples were processed in 1 hour, after being collected from the hospital. The tissues were dissociated with Human Tumor Dissociated kit (Miltenyi Biotec) and filtered using 100µm SmarterStrainers (Miltenyi Biotec). To culture primary cells from patients, we adopted the medium for FTE culturing (Kessler et al., 2015) (41.7ml Advanced DMEM/F12 Medium, 540µl HEPES, 450µl P/S, 4.5µl (10ng/ml) EGF, 2.25ml FBS (5%), 45µl (9µM) Rock inhibitor/Y0503-5MG, Y-27632 dihydrochloride Sigma-Aldrich, termed as FT culturing medium). Immediately after dissociation, cells were transferred into a 6-well plate for primary cell culture with the FT culturing medium at 37°C in a CO_2_ cell incubator, where they were cultured for overnight or long-term (LT, 2 days or 6 days).

For cryopreservation, cells were suspended in 1ml Synth-a-Freeze Cryopreservation Medium (Thermo Fisher). Cell suspension was frozen in freezing containers (Thermo Fisher) with Isopropyl alcohol at −80°C for 24 hours before being transferred into liquid nitrogen. The cryopreservation cells were frozen down immediately after dissociation, stored in liquid nitrogen and recovered in FT culturing medium overnight after thawing. We were not able to generate cDNA from single cells that were immediately thawed from liquid nitrogen. Therefore, the frozen cells were recovered overnight in the FT culturing medium before cell sorting.

#### Cell harvest

For cultured cells, cells were harvested with TrypLE (Thermo Fisher) for 5min at 37°C and dispensed in FACS buffer (1× PBS, 1% BSA, 2mM EDTA, 0.0025% RNasin plus), and then they were processed as for the freshly dissociated cells.

#### Organoid culture

The three-dimensional organoid cultures were conducted as previously described (Kessler et al., 2015). The FT tissue was cut into small pieces of around 2-4 mm in length and incubated in pre-warmed Digestive Medium containing Advanced DMEM/F12 (Gibco), 12mM HEPES (Gibco), 1% Penicillin/Streptomycin (Gibco), 2mg/ml Trypsin (Sigma), 0.5 mg/ml DNase I (Sigma) and 100U/ml Collagenase Type I (Invitrogen); this was incubated for 45min in rotation at 37°C. Using forceps, decellularized tissue was removed and immediately filtered through a 100µm cell strainer and pelleted by centrifugation at 300g for 10min. The cell pellet was washed once with ice-cold DPBS and resuspended in Matrigel Matrix (Corning). Using an 8-well chamber slide (Thistle Scientific), 20µl of Matrigel were placed into each well by drops and the mixture was incubated at 37°C for 20min to allow Matrigel polymerizing and solidifying. Finally, cells were overlaid with pre-warmed organoid medium containing Advanced DMEM/F12, 12mM HEPES, 1% Penecillin/Streptomycin, 1% N2 Supplement (Thermo Fisher), 2% B-27 Supplement (Thermo Fisher), 100ng/ml human Noggin (Peprotech), 100ng/ml human FGF10 (Peprotech), 1mM Nicotinamide (Sigma), 0.5μM TGF-b R Kinase Inhibitor VI (SB431542; ThermoFisher), R-Spondin1 protein (RSPO1) and 25% Wnt3a conditioned medium (ATCC). RSPO1 protein was produced in-house using a HA-RSPO1-Fc 293T cell line (Cultrex). Protein expression was performed according to Culturex protocol while protein purification was done using Pierce Protein A Agarose kit (ThermoFisher Scientific).

#### Cell sorting

To sort freshly dissociated cells or harvested cells, the cells were dispensed in the FACS buffer (1× PBS, 1% BSA, 2mM EDTA, 0.0025% RNasin plus) and stained with EpCAM-APC antibody (Miltenyi Biotec) and CD45-FITC antibody (BioLegend) for 30 min at 4°C in the dark. Afterwards, cells were washed once with FACS buffer and kept on ice in the dark. 10µg/µl DAPI was added 10min before cell sorting with SH800 sorter (Sony). The alive EpCAM+CD45-DAPI-fallopian tube epithelial cells were sorted into 96-well plates with one cell per well. Each batch was sorted with bulk controls (10 or 100 cells) and empty controls. A small number of EpCAM-CD45-DAPI- or EpCAM-CD45+DAPI-cells were sorted as control for epithelial cells. The cells were sorted into 4µl lysis buffer with 0.1µl RNase inhibitor (Clontech), 1.9µl 0.4% Triton X-100, 1µl 10µM 5′-biotinylated oligo-dT_30_VN (IDT) and 1µl 10mM dNTP (Thermo Scientific), snap frozen on dry ice and stored at −80°C before single-cell RNA sequencing.

#### cDNA synthesis and library construction

The SMART-seq2 protocol (Picelli et al., 2014) was used to generate single-cell cDNA and libraries with optimization and automation. To lyse the cells, the aforementioned lysis buffer with single cells were removed from −80°C, heated at 72°C for 3min and then kept at 4°C before adding the reverse transcription master mix. The reverse transcription step exactly followed SMART-seq2 protocol with 5’-biotinylated TSO (Qiagen, primers see Table S9). The first round of 24-cycle PCR started with 10µl reverse transcription product, 0.125 µl 10µM 5’-Biotinylated ISPCR primers (IDT, primers see Table S9), 12.5 µl KAPA HIFI PCR Master Mix (Roche Diagnostics) and 2.375 µl RT-PCR grade water (Invitrogen). PCR products were cleaned up with 0.8:1 Ampure XP beads (Beckman Coulter) with Biomek FxP Laboratory Automation Workstation (Biomek).

The quality of single-cell cDNA was checked on TapeStation with HD5000 Tapes and reagent (Agilent) and single-cell qPCR of *GAPDH* or *ACTB* with QuantiNova SYBR Green PCR Kit (Qiagen, primers see Table S9). The concentration was measured with Quant-iT™ PicoGreen™ dsDNA Assay Kit (Invitrogen) on the CLARIOstar Plate Reader (BMG Labtech). The wells where Ct values of *GAPDH* or *ACTB* were less than 20 were defined as the wells with good-quality cDNA. Good-quality cDNA was cherry-picked and normalized to 2µg/ml with EB (Qiagen) on the Biomek FxP Workstation into a new 384-well plate (4titude).

We used the miniaturized Nextera XT (Illumina) protocol (Mora-Castilla et al., 2016) with Mosquito HTS (ttplabtech) to generate libraries from normalized single-cell cDNA in a 384-well Endure plate (Life Technology) with 400nl input cDNA per reaction. The Nextera index set A and D were used in parallel to multiplex 384 cells per batch. All 384 single-cell libraries in each batch were pooled and cleaned up with 0.6:1 Ampure XP beads (Beckman Coulter). The multiplexed RNA-seq library was normalized to 4nM with Suspension Buffer (provided in Nextera XT kit) to be sequenced on the NextSeq 500 Sequencer (Illumina).

#### Whole-exome sequencing

The buffy samples and tumor biopsies were stored at −80 °C. Tumor biopsies were frozen in the O.C.T. Compound (Fisher Scientific) and sectioned into 1.5mL tubes for DNA extraction. The tumor sites were confirmed by H&E staining. Genomic DNA (gDNA) was extracted from buffy and tumor samples with the DNeasy Blood & Tissue Kits (Qiagen). The whole-exome sequencing libraries were prepared with the SureSelect Low Input Target Enrichment System (Agilent) and sequenced pair-ended on Illumina HiSeq. The reads were filtered with Trim Galore!, and mapped with Bowtie2 (Langmead and Salzberg, 2012). The somatic mutations were called via Platypus (Rimmer et al., 2014) and VarScan2 (Koboldt et al., 2013), and then analyzed by R package VariantAnnotation (Obenchain et al., 2014).

#### Tissue sectioning and Laser Capture Microdissection (LCM)

Tissue sectioning, H&E staining and the assessment by a gynecological oncology pathologist were performed as previously described (Hellner et al., 2016). Tissue biopsies were frozen down with Fisher Healthcare™ Tissue-Plus O.C.T. Compound (NEG-50, Richard-Allan Scientific) and sectioned by 10µm in CryoStar NX50 Cryostat (Thermo Scientific). The tissue sections were stored at −80°C before proceeding with the staining step.

The pre-chemotherapy tumor biopsies were fresh frozen in OCT and sectioned onto polyethylene naphthalate membrane (PEN) glass slides (MembraneSlide 1.0 PEN, Zeiss) at 6µm and immediately stored at −80°C until the laser capture microdissection (LCM). Prior to LCM, the slides were taken out from −80°C, immediately immerged in 70% ethanol for 2 min, in Cresyl Violet (Sigma Aldrich) for 2 min, rinsed in 95% ethanol five times to remove residual cresyl violet and dried at room temperature for 2min.

LCM was conducted on the PALM Laser Microdissection System (Zeiss) and the captured cells were collected onto the adhesive caps of PCR tubes (AdhesiveCap 500 opaque, Zeiss), then immediately resuspended in 10µl lysis buffer (RNAqueous™-Micro Total RNA Isolation Kit, Invitrogen). RNA was extracted according to the manufacturer’s instructions and 3µl of eluate before DNase treatment wass kept for Whole Genome Amplification and stored at −20°C. The remaining RNA was DNase treated, assessed with the Agilent TapeStation 2200 (Agilent) and then stored at −80°C.

#### Bulk RNA sequencing

The SMARTer Stranded Total RNA-Seq kit v2 - Pico Input (Takara) was used to prepare sequencing libraries which were indexed, enriched by 15 cycles of amplification, assessed using the Agilent TapeStation and then quantified by Qubit. Multiplexed library pools were quantified with the KAPA Library Quantification Kit (KK4835) and sequenced using 75bp pair end reads on the Illumina NextSeq500 platform. FastQ files were trimmed for adapter sequences and quality. Trimmed reads were then mapped to the human hg19 reference genome using STAR (v2.4.2a) and read counts were obtained using subread FeatureCounts (v1.4.5-p1).

#### Immunofluorescent staining

To do immunofluorescent staining on the frozen OCT-embedded tissue sections, slides were taken out from −80°C and thawed at room temperature for 5 min. The slides were washed with ice-cold PBS twice to remove the OCT and then fixed in ice-cold 4% PFA for 10min. The fixed slides were washed with PBS/Glycine buffer to remove residual PFA and then permeabilized with Permeabilization Buffer (1×PBS, 0.5% Triton X-100) for 10min at RT. A hydrophobic circle was drawn around the tissue region with the PAP pen (Abcam) and dried for 5min at room temperature (RT). The permeabilized slides were washed in Wash Buffer (1×PBS, 0.2% Triton X-100, 0.05% Tween-20) and blocked in Blocking Buffer (Wash buffer, 10% donkey serum, Sigma) for 1-2 hours. The diluted primary antibodies were incubated at 4°C overnight or at RT for 1-2 hours (antibodies see Table S9). The slides were washed and incubated with the secondary antibodies (1:200) and DAPI (1:1000) at RT for 1-2 hours. The stained slides were washed, mounted with the mounting medium without DAPI (Vector Laboratories) and covered. Then they were dried for 1-2 hours at RT in dark before being imaged using a Confocal Microscope (Zeiss 780).

#### Immunohistochemistry

The Formalin-Fixed Paraffin-Embedded (FFPE) samples were sectioned into 2.5 or 4 µm. We used the Leica Bond Max autostainer (Leica Microsystems) or Autostainer plus Link 48 (Dako) to automatically stain the tissue sections. The standard Heat Induced Epitope Retrieval (HIER) in retrieval buffer pH 6 was used. Tissue sections were kept at 100 °C for 20min and incubated with the primary antibody or lgG control for up to 1 hour (antibodies see Table S9). We used the Aperio slide scanner (Aperio) to scan the stained slides at 20× or 40× magnification.

### QUANTIFICATION AND STATISTICAL ANALYSIS

#### Preprocessing and quality control of scRNA-seq data

Sequencing files were trimmed by Trim Galore!, mapped to UCSC hg19 human genome assembly by STAR (Dobin et al., 2012) and counted by FeatureCount (Liao et al., 2014). The quality of the expression data was checked by the R package scater (McCarthy et al., 2017) and the low-quality cells were filtered out from further analysis. We used three criteria to filter cells for further analysis: first, the number of genes detected per cells was more than 1200 and less than 7500; second, the total number of read counts was more than 0.1 million per cell; third, the percentage of counts of the top highly expressed 200 genes was less than 85% to ensure enough complexity of the transcriptomes. The upper cutoff for numbers of genes per cells was aimed to filter out the putative doublets. After filtering, the numbers of detected genes per cell were between 1,202 and 7,496 (mean: 3,665). The numbers of total reads per cell were between 102,319 and 9,668,129 (mean: 763,233).

#### Detecting doublets

To effectively detect doublets, we used R package DoubletFinder (McGinnis et al., 2019). In brief, we combined all fresh cells (secretory cells, ciliated cells, fibroblasts and leukocytes), constructed a Seurat S4 object (Butler et al., 2018) and ran DoubletFinder with its default pipeline (PCs = 1:10, pN = 0.25, pK = 0.07).

#### Visualization of expression data

The uniform manifold approximation and projection (UMAP) plots were based on the functions of runUMAP and plotUMAP, and the violin plots of expression were plotted by the function plotExpression from the R package scater. Heatmaps were plotted by R packages pheatmap (Kolde, 2012) and ggplot2 (Ginestet, 2011).

#### Comparing fresh and cultured cells

To investigate the effects of cell culture, we first combined secretory cells from Patient 11553 and Patient 15072. Then we used limma-voom to identify the DE genes in three pairs of comparisons, namely fresh versus overnight, fresh versus LT and overnight versus LT. The threshold of DE genes was fold-change ≥ 2 and FDR < 0.05. The total number of DE genes affected by culture condition was 4061. The GO analysis was conducted by Gorilla with cutoffs at FDR < 0.05.

#### Pseudotime analysis

We used the R package PhenoPath (Campbell and Yau, 2018) to perform the pseudotime analysis that projected the high-dimensional transcriptomic data to one dimension, in which we compared fresh, overnight cultured and long-term cultured cells. The expression matrix that was normalized by the R package scater was used as the input and the first principal component was used as the initialization of the latent trajectory to accelerate the convergence.

#### Filtering to select fresh secretory cells

Only the cells that belonged to the secretory cluster and the fresh group were kept expect for the cells from the 15066-left tube (15066L), because the cells from 15066L formed a patient-specific cluster (Figure 1F) and it was reported to have salpingitis on this side. Secretory cells with strong expression of *KRT7* and *EPCAM* (log [CPM+1] > 2), no expression for *PTPRC* (log [CPM+1] = 0) or *CCDC17* (log [CPM+1] < 1) and no observed copy number variants were kept for the clustering analysis. Copy number variants were detected by HoneyBADGER with default settings (Fan et al., 2018).

Single-nucleotide variants (SNVs) were called from piled up BAM files by MonoVar (Li et al., 2009; Zafar et al., 2016) with the recommended parameters (samtools mpileup -BQ0 - d10000 -f ref.fa -q 40 -b filenames.txt | monovar.py -p 0.002 -a 0.2 -t 0.05 -m 2 -f ref.fa -b filenames.txt -o output.vcf). SNVs in exome regions were intersected by the Bedtools (Quinlan and Hall, 2010).

#### Differentially-expressed-based clustering approach

We used an in-house clustering algorithm to identify the cell populations across patients. In brief, the clustering method contained three steps: (1) an initial clustering step within the individual patients by Spectral Clustering (Karatzoglou et al., 2004; Ng et al., 2002); (2) calculation of distance between each pair of initial clusters by differential expression (DE) analysis; (3) a final hierarchical clustering across different patients based on the calculated distance matrix (Figure 1B).

The clustering approach for clinical scRNA-seq data required the input of an expression matrix, *M* (*m × n*), where the *m* rows corresponded to genes/features and the *n* columns corresponded to cells. The patient source of all cells was denoted by a vector, *p*. The length of *p* was equal to number of cells (*n*) and the number of different values in *p* was equal to the number of patients (*k*). For the dataset containing *k* patients (*k* > 1), our method consisted of three main steps.

The count matrix was preprocessed by log-normalization, selection of high variance genes (Satija et al., 2015), centered and dimensionally reduced by the principal component analysis. The first step was initial clustering of the cells within individual patients by the weighted k-nearest neighbors spectral clustering provided by kknn (Hechenbichler and Schliep, 2004). After spectral clustering, each cell was assigned to a patient-specific initial cluster. Cells from different patients were clustered into separate patient-specific clusters.

The second step was pairwise differential expression (DE) analysis by limma voom (Law et al., 2014; Phipson et al., 2016; Ritchie et al., 2015) between initial clusters across patients to build a distance matrix, because it was reported to be one of the top ranking DE analysis methods for scRNA-seq (Soneson and Robinson, 2018). There were two modes to calculate the distance. The first one was unweighted, meaning that the distance between two initial clusters was defined as the number of differentially expressed genes (DEGs) between them. The second mode was weighted. The DE genes had different weights in the distance. The DE genes with higher fold change and larger ratio of expression had higher weight. The weighted method had advantages in identifying rare but functionally distinguished cell types. As a result, a distance matrix of functional dissimilarity was constructed among all initial clusters.

In the last step, all initial clusters were assigned to the final clusters by hierarchical clustering based on the aforementioned distance matrix. Accordingly, the single cells were assigned to the final cluster containing the initial cluster that the cells belong to. The hierarchical tree was plotted to help dividing the final clusters.

#### Identifying marker genes

The identification of markers was based on limma-voom, which has good performance with scRNA-seq data (Soneson and Robinson, 2018). To spot the positive markers in the target cluster, we conducted the differential expression analysis between the target cluster and background (non-target) clusters. Only if one gene emerged in all the pairwise comparisons between the target cluster and background clusters with FDR < 0.05 and log_2_ fold-change > 0.4, this gene was defined as the marker gene of this target cluster.

#### Gene ontology enrichment analysis

Gene ontology enrichment analysis was conducted on GOrilla (last updated on May 26, 2018) (Eden et al., 2007; 2009). The statistical overrepresentation was conducted by the PANTHER Gene list analysis (Thomas, 2003). Pathway enrichment analysis was done by the DAVID Bioinformatics Resources 6.8 (Huang et al., 2009b; 2009a). The threshold of significance was false discovery rate (FDR) < 0.05.

#### Bulk expression data

TCGA data was downloaded from the UCSC Xena Data Hub (https://tcga.xenahubs.net) (Goldman et al., 2018) and the Broad firehose (https://gdac.broadinstitute.org), which was derived from the TCGA Data Coordinating Centre (version: Jan 2016). We used the “IlluminaHiSeq UNC” RNA-seq dataset (version: 2017-10-13) and the “TCGA OV gene expression subtype” phenotype data (version: 2016-05-27). The AOCS dataset was downloaded from GSE9899. The other microarray datasets were retrieved from the R package CuratedOvarianData (Ganzfried et al., 2013).

#### Deconvolution of bulk expression data

On the basis of our FTE scRNA-seq data, we firstly selected the marker genes for five FTE subtypes, namely Differentiated (C3), KRT17 cluster (C4), Cell cycle (C9), EMT (C7) and Ciliated signatures by using the following cutoffs: (1) log_2_ fold-change ≥ 1.5; (2) FDR ≤ 0.01; (3) dispersion ≥ 0.2 that was measured by the variable loadings in the principal component analysis of TCGA RNA-seq data; (4) the correlation with each other markers was less than 0.9 in TCGA data to avoid redundancy. We calculated the reference matrix (Figure 4A) by BSEQ-sc (Baron et al., 2016) that measured that average expression levels of marker genes in each cell subtype. We used this reference matrix to deconvolute the RNA-seq data from TCGA, the microarray data from AOCS and CuratedOvarianData and the OXO-PCR RNA-seq data by CIBERSORT as previously described (Newman et al., 2015), which is based on the u-parameter support vector regression and is robust to noise and outliers. The CIBERSORT was ran in the relative mode with non-logged expression matrices. For the microarray data, if a gene corresponded to multiple probes, we selected the probe that had the highest average expression level to represent that gene. The deconvolution analysis generated scores of the five transcriptomic signatures in each tumor sample (Figure 4A). This score of a signature can be interpreted as the proportion of the corresponding cell state in a tumor sample.

#### Survival analysis

The survival analysis was conducted by fitting a Cox proportional hazards regression model. This analysis used age, grade and residual disease as covariates. We repeated the survival analysis in TCGA RNA-seq dataset, AOCS microarray dataset and other seven microarray datasets from the CuratedOvarianData database (Ganzfried et al., 2013). The criterion of selecting datasets was that a dataset contained over one hundred clinical samples. For the datasets from CuratedOvarianData that contained both low and high grades or multiple histological types, we used the multivariate analysis to eliminate the effects of confounding factors. In the multi-variate survival analysis, we dichotomized grades to low-grade (Grade 1) and high-grade (Grades 2 and 3) and dichotomized histological subtypes to serous and non-serous (endometrioid, clear cell, mucinous, undifferentiated, mixed and *etc.*) (Table S8). EMT scores and overall survival were negatively correlated in the independent eight datasets (P < 0.05), except for the “GSE32062.GPL6480” dataset (see Table 1). The hazard ratios of EMT scores in all the nine independent datasets were larger than 1 (range: 1.88 – 3.31). For the survival curve of the meta-analysis (Figure 5C), we dichotomized the EMT-low and EMT-high groups by the median of EMT scores of each dataset to avoid batch effects.

## Acknowledgments

We thank the WIMM Sorting Facility, the WIMM Single Cell Facility and the Computational Biology Research Group (CBRG) for their help in this study. We thank Leticia Campo for technical help. This work was supported by Ovarian Cancer Action and Oxford Biomedical Research Centre, National Institute of Health Research. CY was supported by a UK Medical Research Council Research Grants (MR/L001411/1, MR/P02646X/1), a Wellcome Trust Core Award Grant Number 090532/Z/09/Z and by The Alan Turing Institute under the EPSRC grant EP/N510129/1.

---

**Supplementary Figure 1.**
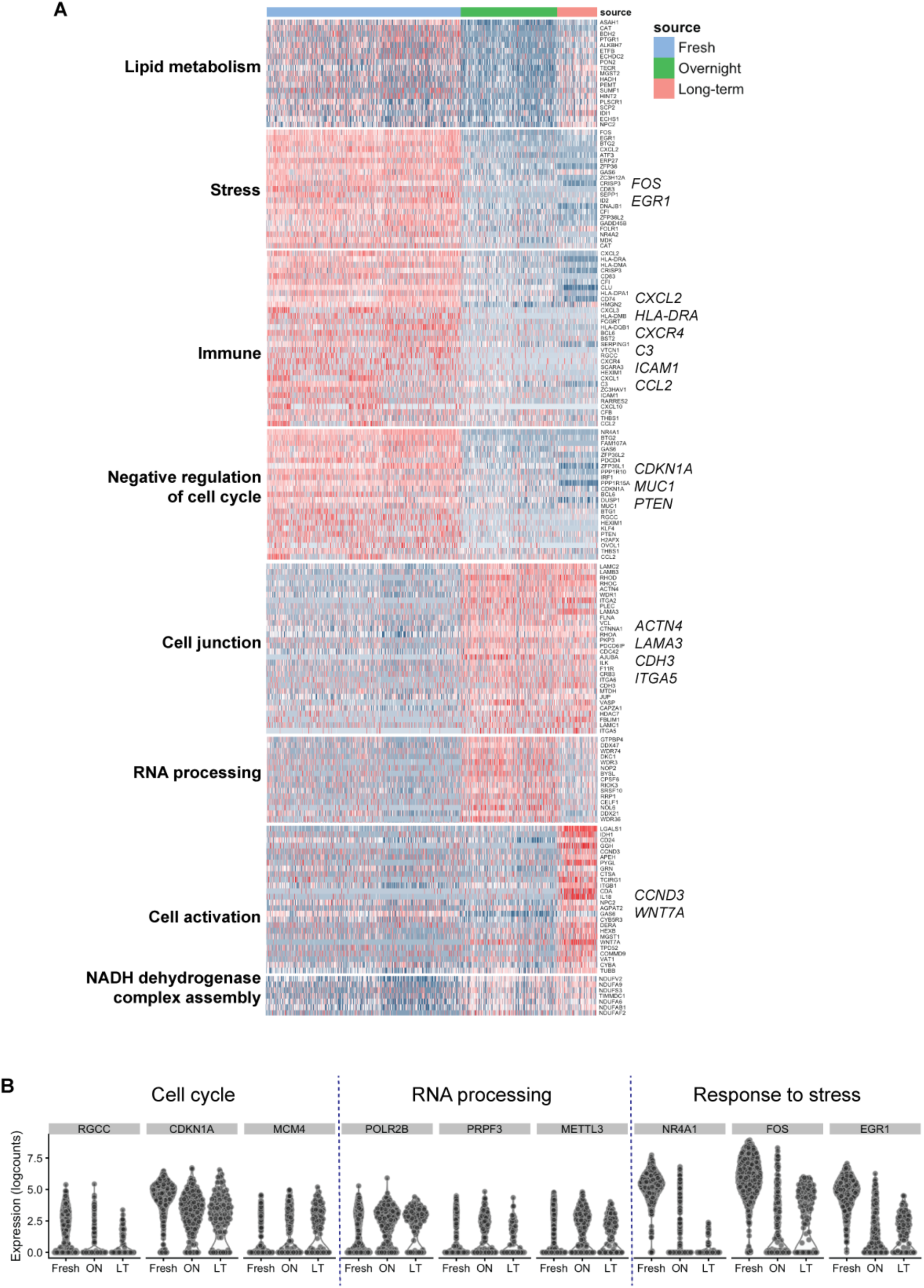
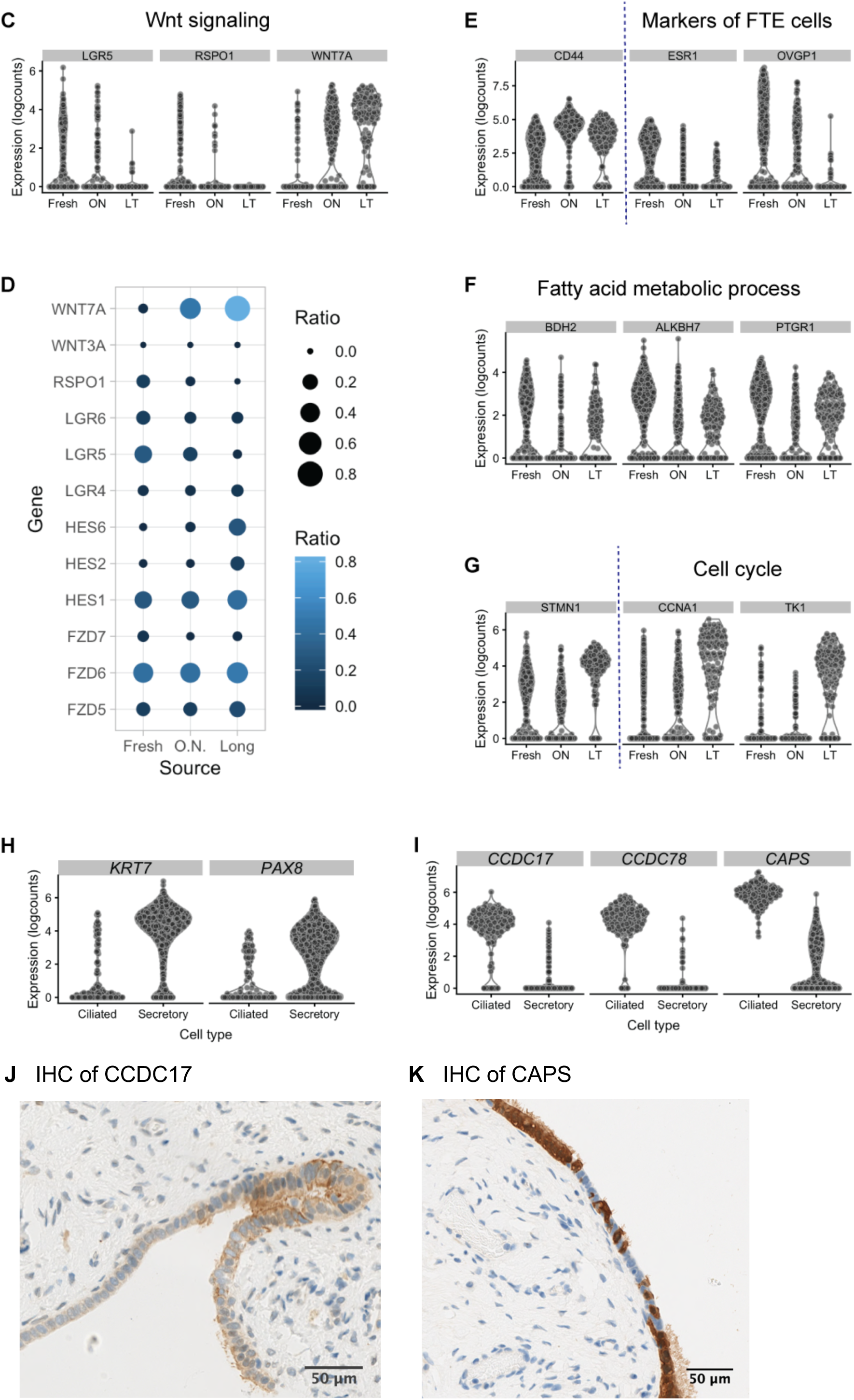
Landscape of single-cell transcriptomes in fallopian tube epithelium. Cryopreservation and cell culture are common techniques to maintain alive primary cells, especially for rare clinical samples. We tested the effect of the cell maintenance approach with primary cells from human fallopian tubes. We applied the pseudotime analysis (Campbell and Yau, 2018) on the transcriptome of primary cells under three conditions (freshly dissociated, overnight cultured and long-term cultured) from Patients 11553 and 15072, because they had cells from all three conditions. The cultured cells included short-term overnight culture and long-term culture (2-day or 6-day). Based on the predicted pseudotime of each cell, fresh cells were more similar to the long-term (LT) group compared to the overnight group (see also Figure 1E), possibly because some pathways were transiently switched on or off in the overnight group, such as the fatty acid metabolic process and the RNA processing pathway. After cryopreservation, ciliated cells raised the expression of genes related to regulation of cell cycle but downregulated the genes related to cilium organization (see also Table S2 and Table S3). This suggests that the maintenance of ciliated features may depend on the signaling from the external environment, which agrees with the previous studies that estrogen can affect ciliogenesis in fallopian tubes (Nayak et al., 1976) and that ciliated cells are not preserved in culturing conditions (Karst and Drapkin, 2012). (A) Heatmap shows the differences among fresh, overnight-cultured and LT-cultured groups. Rows represent differentially expressed genes ordered by the gene ontology that they belong to. Columns represent single cells with the annotation bar that indicates the condition of each cell. Compared to fresh cells, the overnight cultured cells were upregulated in genes (fold-change > 2, FDR < 0.05) enriched in cell division/cell cycle, NADH dehydrogenase complex assembly, cell junction assembly, DNA-templated transcription/RNA processing and *etc.* The downregulated genes were enriched in cell communication, immune process, response to stress, regulation of cell motility and negative regulation of cell cycle (e.g. p21/*CDKN1A*). Hence, the culture condition impacts cell metabolism, cycling and signaling. (B) Violin plots show representative genes in three pathways that are differentially expressed between fresh and overnight-cultured groups. FDR < 0.05, by limma voom. Overnight culture induced profound differential expression changes in pathways related to cell cycle (e.g. *RGCC*, p21 and *MCM4*), RNA processing (e.g. *POLR2B*, *PRPF3* and *METTL3*) and stress response (e.g. *NR4A1*, *FOS* and *EGR1*) (see also Table S2). (C) Violin plots show the expression of representative genes in the Wnt signaling pathway that is dysregulated by the culturing conditions. FDR < 0.05, by limma voom. (D) Dot plots show that the Wnt and Notch signaling pathways are influenced by culture condition. The culture condition changes the proportion of cells that express genes related to the Wnt pathway. The size and color of each dot represents the proportion of cells that express genes (rows) across three conditions (columns) as shown in the scales. The Wnt and Notch signaling pathways are essential in regulating the growth of fallopian tube organoids (Kessler et al., 2015) and the maintenance of stem cells. Our analysis indicates that the Wnt and Notch signaling pathways are affected by the culture condition, which may not represent the in-situ status. (E) Violin plots show the dysregulated expression of three genes (*CD44*, *ESR1* and *OVGP1*) after overnight culture. FDR < 0.05, by limma voom. (F) Violin plots show that genes that are enriched in fatty acid processing are transiently switched off after overnight culturing. FDR < 0.05, by limma voom. (G) Violin plots show that three genes (*STMN1*, *CCNA1*and *TK1*) related to the cell cycle pathway are significantly upregulated and expressed in most of cells after LT culturing. FDR < 0.05, by limma voom. (H) Violin plots show the specific expression of secretory markers (*KRT7* and *PAX8*) in fresh secretory cells compared to fresh ciliated cells. (I) Violin plots show the specific expression of ciliated marker genes (*CCDC17*, *CCDC78* and *CAPS*) in fresh ciliated cells compared to fresh secretory cells. (J) Immunohistochemistry (IHC) staining of a ciliated cell marker CCDC17 in the Formaldehyde Fixed-Paraffin Embedded (FFPE) human fallopian tube section. This CCDC17 antibody stained the cytoplasmic and membrane region of ciliated cells brown. (K) IHC staining of a ciliated cell marker CAPS in the FFPE human fallopian tube section. The ciliated cells show strong brown staining of CAPS in the cytoplasm, which is absent in the secretory cells.

**Supplementary Figure 2.**
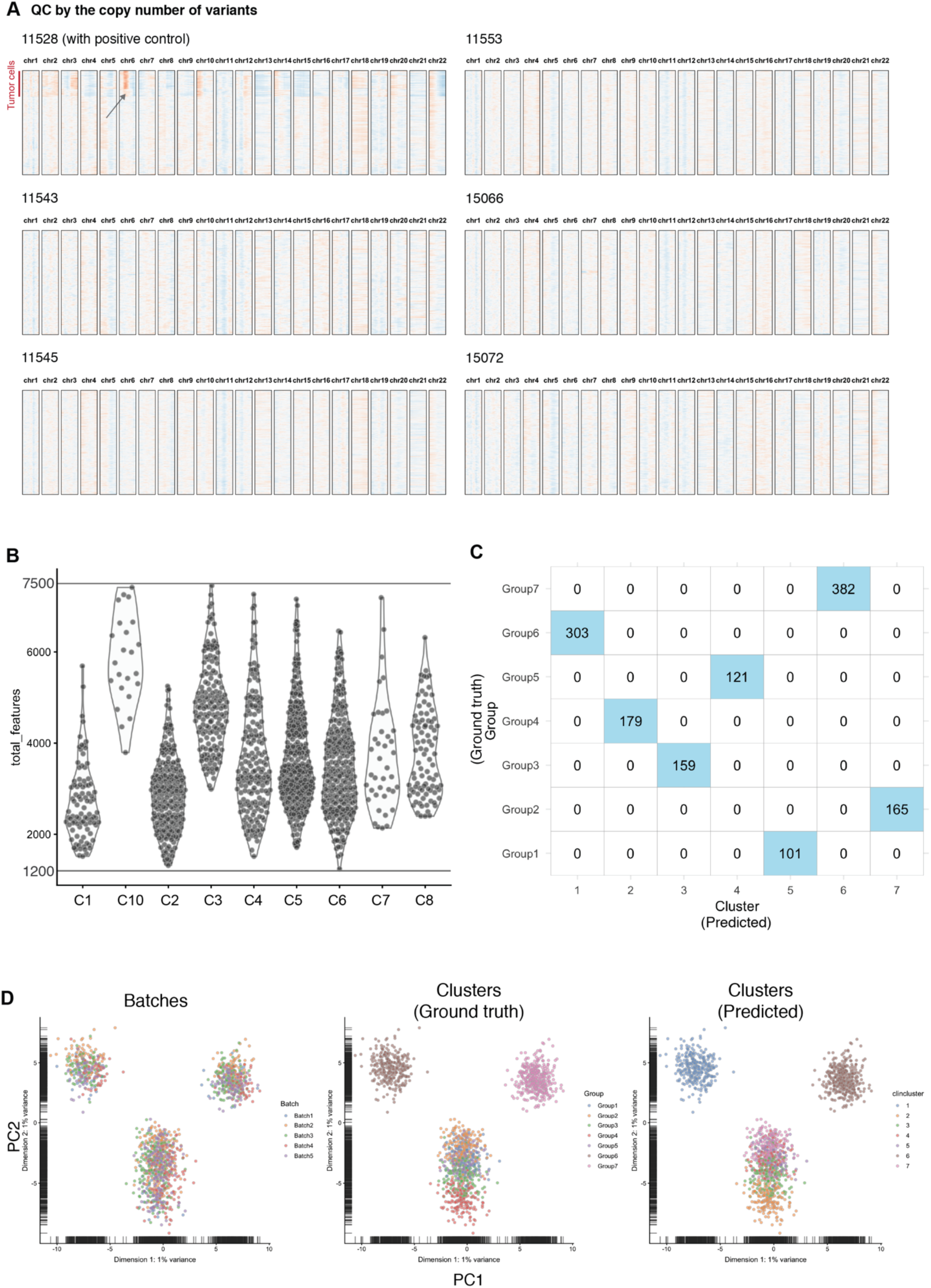
Four novel secretory subtypes and their molecular features. (A) Relative copy number variation inferred from the expression data. Note that only a small proportion of cells in patient 11528 seemed to have copy number variation at Chr6 (arrow). These cells also showed a mutation of TP53 in the single-cell RNA-seq data. Each heatmap contains the cells from one patient. In the heatmap, each row is a single cell and columns represent the positions on chromosomes. Each block represents one chromosome in the order from Chr1 to Chr22. The color scale denotes the possibility of amplification or deletion. Red means amplification and blue means deletion as shown in the scale bar. In the left tube from Patient 11528, we found that a TP53-mutant FTE cluster with distinctive expression profile shared the same TP53 somatic mutation (chr17: 7577515T>G) with the tumor sample. This somatic mutation induced an increased expression level of TP53 in this cluster. Further analysis revealed that this cell subpopulation shared the same amplification on Chromosome 6 and deletion on Chromosome 13, 15 and 17 with tumor samples. It indicates that these TP53-mutant cells may be prelesion or metastasized tumor cells. (B) Violin plot shows that all cells that underwent the clustering analysis and passed the filter of doublets (number of genes detected ≤ 7,500). The C7 (EMT) cluster has a similar distribution of transcriptomic complexity as other clusters (C4, C5 and C8). Two quiescent populations (C1 and C2) and the stress cluster (C6) have lower numbers of genes detected per cell. The cell cycle cluster (C9) and the differentiated cluster (C3) have higher numbers of genes detected. (C) Heatmap shows the comparison between the clustering results (column) and the ground truth of populations (rows) of the simulated dataset. The simulated dataset contains 1,410 cells distributed across 5 batches and 7 populations. To verify that our in-house differential-expression-based (DE-based) clustering is capable of identifying cell populations precisely, we tested our clustering method with a simulated data generated by Splatter (Zappia et al., 2017). This simulated dataset has the same sample size as our dataset of fresh secretory cells. The numbers of batches and populations are consistent with the ones of our dataset. We preformed the DE-based clustering method on the simulated dataset and found that the clustering result completely matches the ground truth (adjusted rand index = 1, see also Figure S3D). The result demonstrates that our approach can precisely cluster data with confounding batch effects. (D) Three PCA plots respectively show the distribution of simulated cells colored by the batches/patients (left panel), ground truth of populations (middle panel) and the predicted results by clustering (right panel).

**Supplementary Figure 3.**
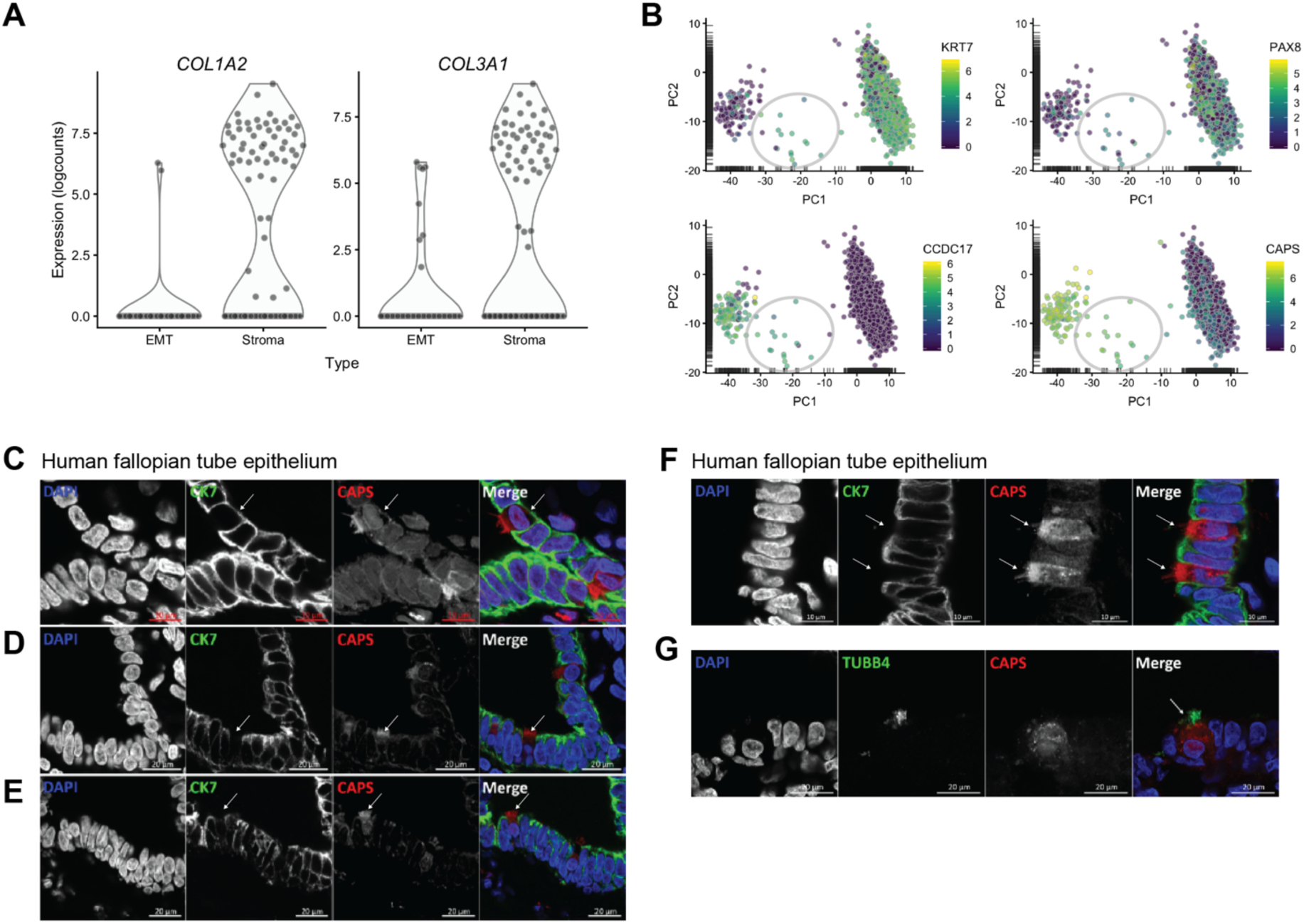
The non-traditional cell subtype in the human fallopian tube epithelium. (A) Violin plots show that two stromal markers were specifically expressed in mesenchymal cells but not in the EMT cluster of FTE secretory cells. (B) PCA plots show that the intermediate cell population (grey circle) has the expression of both secretory markers (*KRT7* and *PAX8*) and ciliated markers (*CCDC17* and *CAPS*). (C-F) IF staining of CK7 (KRT7) and CAPS in human fallopian tube sections shows that secretory cells are CK7 positive and CAPS negative (arrows). (G) IF staining of both TUBB4 and CAPS in human fallopian tube sections shows that a TUBB4 positive ciliated cell is also CAPS positive (arrow). It demonstrates that CAPS is a marker of ciliated cells (see also Figure S1K).

**Supplementary Figure 4.**
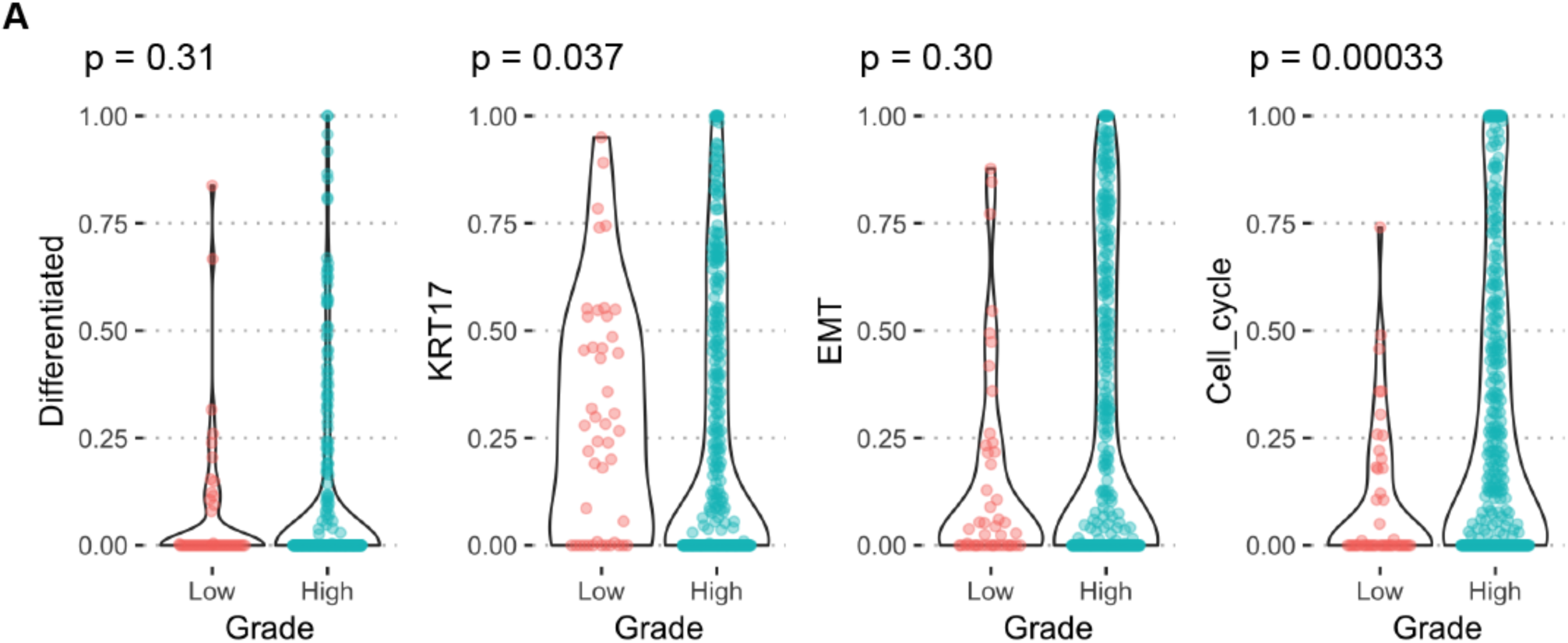
The repertoire of phenotypic heterogeneity of serous ovarian cancer revealed by deconvolution analysis. Violin plots visualize the different composition of cell states between low-grade and high-grade serous ovarian cancer in the AOCS dataset and GSE1726. P-values were calculated by the Wilcoxon test between the low-grade and high-grade tumors. (See also Figure 4F)

**Supplementary Figure 5.**
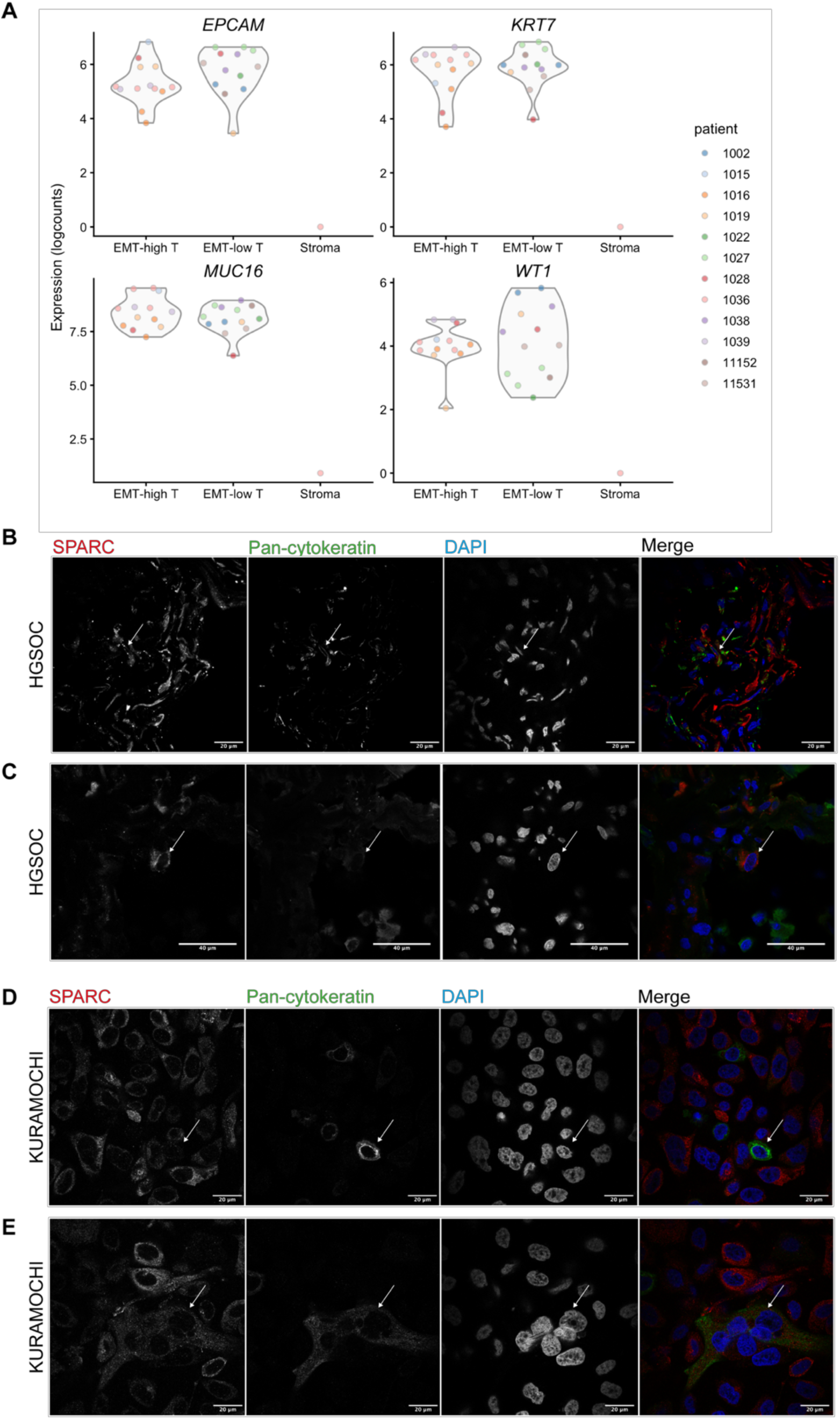
Mesenchymal-like tumor cells in pre-chemotherapy tumors and KURAMOCHI cell line. (A) Expression levels of ovarian cancer markers (*EPCAM*, *PAX8*, *MUC16*, *WT1*) were significantly higher in both EMT-high and EMT-low tumor samples compared to stromal samples in which these markers showed almost no expression. (B) The SPARC and pan-cytokeratin double positive cells most probably represent the EMT or MET tumor cell population (see also Figure 6C). (C) IF staining shows a SPARC positive and pan-cytokeratin negative population with similar nuclear sizes as tumor cells implying that it is a mesenchymal-like tumor cell (see also Figure 5D). (D) KURAMOCHI cells exhibited two phenotypes, a SAPRC+ mesenchymal-like phenotype and a pan-cytokeratin+ epithelial phenotype (arrow). (See also Figure 5E) (E) A SPARC and pan-cytokeratin double positive population also exists in KURAMOCHI cells (arrow).

